# Size-dependent membrane remodelling at Myomerger clusters during myoblast fusion

**DOI:** 10.64898/2026.07.31.741798

**Authors:** Upasana Mukhopadhyay, Madhura Chakraborty, Bidisha Sinha

## Abstract

Myomerger drives fusion pore formation after hemifusion of membranes during myogenic cell-cell fusion in myogenesis, but its mechanical impact has been characterised largely in liposomes. In differentiating C2C12 cells, hemifusion began at 8 hr and peaked at 24 hr, whereas the earliest content-mixing events appeared only at 24 hr - Myomerger is therefore present much before it is competent to complete fusion. Using Interference Reflection Microscopy (IRM), we found that basal membrane tension in hemifusing cells was indistinguishable from that of non-fusing cells at 8-24 hr, and became significantly higher by 48 hr. Knockdown showed that Myomerger switched across this window from tension-reducing to enhancing cell-scale tension, unlike Myomaker, which enhanced tension in both the early and late phases. Addressing the local impact, Stimulated Emission Depletion microscopy (STED) imaging resolved Myomerger clusters built from a ∼40 nm basic unit that aggregate further. Small clusters induced nanometric outward bulges, a topology rare outside Myomerger-enriched regions; these bulges damped membrane fluctuations and showed higher lipid compaction at the centre, as deciphered by Flipper-TR FLIM, at the lateral as well as the basal membrane, and may be central to the hemifusion-to-pore transition. Larger clusters indented the membrane, showed enhanced fluctuations and associated with the clathrin-coated pits, consistent with surface clearance by endocytosis. Myomerger colocalized poorly with actin fibres and cholesterol-enriched nanodmains despite lying close to both. Together, these findings establish size-based clustering of Myomerger as a determinant of membrane topology and lipid compaction when hemifusion peaks and content mixing begins.

## Introduction

Myotubes – fundamental units of skeletal muscle - are formed and sustained by cell-cell fusion of myoblasts through the process of myogenesis. A critical step in myogenesis – especially during the fusion of myoblasts – is the requirement to overcome thermodynamic barriers that restrict cellular membranes to merge. Although the mechanism of myoblast fusion has been extensively studied in the Drosophila model, the fusion of cells in vertebrate model systems – from mice to human – has been shown to involve different set of fusogens – Myomaker and Myomerger. These transmembrane proteins are necessary and sufficient for fusion(Chen et al., 2020; Leikina et al., 2018) and hence under stringent control in cells. Myomaker and Myomerger are believed to function independently to regulate distinct steps of fusion. Myomaker, a muscle-specific transmembrane protein expressed on both fusing myoblast membranes, is believed to control the early hemi-fusion stage. In this stage, the membranes of two cells merge without mixing their cytoplasmic contents via pore formation (Millay et al., 2013). Conversely, Myomerger is believed to act primarily as a membrane stressor on the hemi-fused membrane to aid the process of pore formation, thereby completing fusion (Golani et al., 2021). The 221 amino acid-long Myomaker is required on both membranes to sustain the hemi-fusion step, whereas the smaller 84 amino acid Myomerger only needs to be present on one of the fusing membranes to initiate pore formation and finalize fusion(Quinn et al., 2017). In vertebrate myoblast fusion, experimental evidence indicates that Myomerger applies mechanical stress to the membrane to facilitate pore formation. Notably, when Myomerger knockout myoblasts—which normally fail to fuse—are subjected to external mechanical stress, such as hypotonic shock or other membrane-stressing agents, recovery in their fusion rate is observed (Leikina et al., 2018). This strongly suggests a critical link between membrane tension and fusion. Literature also suggests that Myomerger can introduce positive curvature to outer leaflet in artificial lipid bilayers (Gamage et al., 2022; Golani et al., 2021) however, there is little understanding of how Myomerger affects cell membranes. It is well established in insects that cortical tension increases during myoblast fusion (Kim et al., 2015; Kozlov & Chernomordik, 2015). Theoretical models have proposed that lateral membrane tension can play a significant role in fusion (Chizmadzhev et al., 2000; Kozlov & Chernomordik, 2015), and recent updates show that early low basal membrane tension coupled with high surface expression of Myomerger is essential to successful fusion (Chakraborty et al., 2022). However, the specific role and the mechanism underlying the stringent basal membrane effective tension remains unclear. Literature also underscores the transient nature of hemi-fusion with a lifetime of ∼15 mins and a basal level of hemi-fusion occurring even in the absence of Myomerger (Golani et al., 2021). Therefore, the role of Myomerger in an early hemi-fusing population is relatively less addressed. However, it remains plausible that Myomerger affects the hemi-fusing cells through modulation of basal membrane tension profile as well as locally aiding in membrane destabilization.

To understand how Myomerger may act as a membrane stressor, it is important to understand the basal mechanical state of membranes and how any change may be measured. Cell membranes have incessant shape fluctuations at microscopic length scales owing to the thermal background and cellular activities. Previous studies have shown that various cellular activities—including ATP-driven processes, membrane cortex dynamics, endo/exocytosis, and protein-drug interactions—affect these fluctuations (Turlier & Betz, 2025)(Biswas et al., 2017). Using the non-invasive imaging method of Interference Reflection Microscopy (IRM) (Biswas et al., 2017, 2019), the spatiotemporal characteristics of membrane fluctuations of the basal plasma membrane of cells can be imaged. Height fluctuations along the vertical (z) axis, can be tracked at high z resolution and diffraction-limited resolution in the horizontal (xy) plane (Biswas et al., 2017, 2019) and can be used to estimate effective mechanical parameters of the membrane. The state of fluctuations and the effective membrane tension derived from the membrane-height time series could provide a deeper understanding of how the protein may affect the membrane. Quantifying Fluctuation amplitude and effective tension would provide information about the global cellular mechanical state as well as map out the spatial distribution. The effective tension in such a case – derived from observing spontaneous fluctuations - is termed as the fluctuation tension ((Biswas et al., 2017)) and must be interpreted distinctly from other measurements – like apparent tension by optical-trap-based measurements – although the two has been shown to be correlated(Ghosh et al., 2025). It must also be noted that the effective tension is derived not solely from passive thermal fluctuations but fluctuations that also have contributions from all players including the cytoskeleton. Furthermore, IRM – coupled to TIRF microscopy, can provide nanometric local topology of microscopic regions where the fusogen Myomeger may be located as a cluster. To validate our observations of basal membrane mechanics and to check the robustness of the membrane mechanical regulations at various heights (z dimension) from the substrate, Fluorescence Lifetime Imaging Microscopy (FLIM) based quantitative and qualitative estimates of lipid compactness of the PM under different physiological conditions were performed. The Fluorescence lifetime of Flipper-TR probe intercalated in a lipid bilayer, is a function of its conformation which in turn depend on the lipid ordering of the bilayer. In cells and multi-phased GUVs, membrane effective tension is known to enhance lifetime due to tension induced lipid ordering and domain coalescence (Lüchtefeld et al., 2024). Flipper-TR lifetime is therefore an indirect measure to non-invasively probe the membrane mechanics of the entire cell.(Colom et al., 2018).

In this work, we have affirmed that Myomerger has a mechano-regulatory role in when hemi-fusion peaks. We have attempted to decipher the underscoring mechanism that helps Myomerger to increase locally the amplitude of membrane fluctuations (SD_time_) in the hemi-fusing population. Our findings reveal that Myomerger clustering aids membrane bending and also aids fusion.

## Results

### Hemifusion starts early with cells progressively getting more tensed than non-fused cells

Since earlier studies have pointed to an early role of Myomerger in myogenesis (Chakraborty et al., 2022) while literature strongly suggests Myomerger is necessary to proceed from hemifused state to the fused state, in this section we sought to understand how early hemifusion events originate, fusion pore starts forming and how the hemifused cells mechanically vary from the rest. We used C2C12 mouse myoblasts for this study. It is to be noted that administering differentiation media (DM) to C2C12 cells generally initiates new fusion events after 48 – 96 hr of DM addition– with fusion events characterized by IRM mages as well as MyHC labelling and multi-nucleated state (Chakraborty et al., 2022). As per literature, it is also known that there is no appreciable change in gene expression profile in myoblasts by 2 hr of DM addition (Yoshida et al., 1998). Hence, the timepoints chosen were from 2 hr – 48 hr for this section. To identify and count the hemi-fusing cells, cells were dual-labelled with lipid probe (red) and content probe (green) and mixed with an unlabeled population (Leikina et al., 2018) (**Fig. 1a**). Fusion was induced in this mixed population using DM. The originally unlabeled cells are expected to acquire only the lipid probe if and only if they hemifused with a neighbouring dual-labelled partner. Such cells were easily distinguished in the sample with red label in their interior and a dual-labelled cell as a neighbour (**Fig. 1b** – white arrows, insets). We observed that such hemi-fusion events start as early as 8 hr post DM addition (∼5%) and increase to ∼ 10% post 24 hr of initiation of differentiation (**Fig. 1c**). By the 48 hr time point, fusion events were observed while the primary hemi-fusing population reduced (**Fig. 1c**).

**Figure 1.**
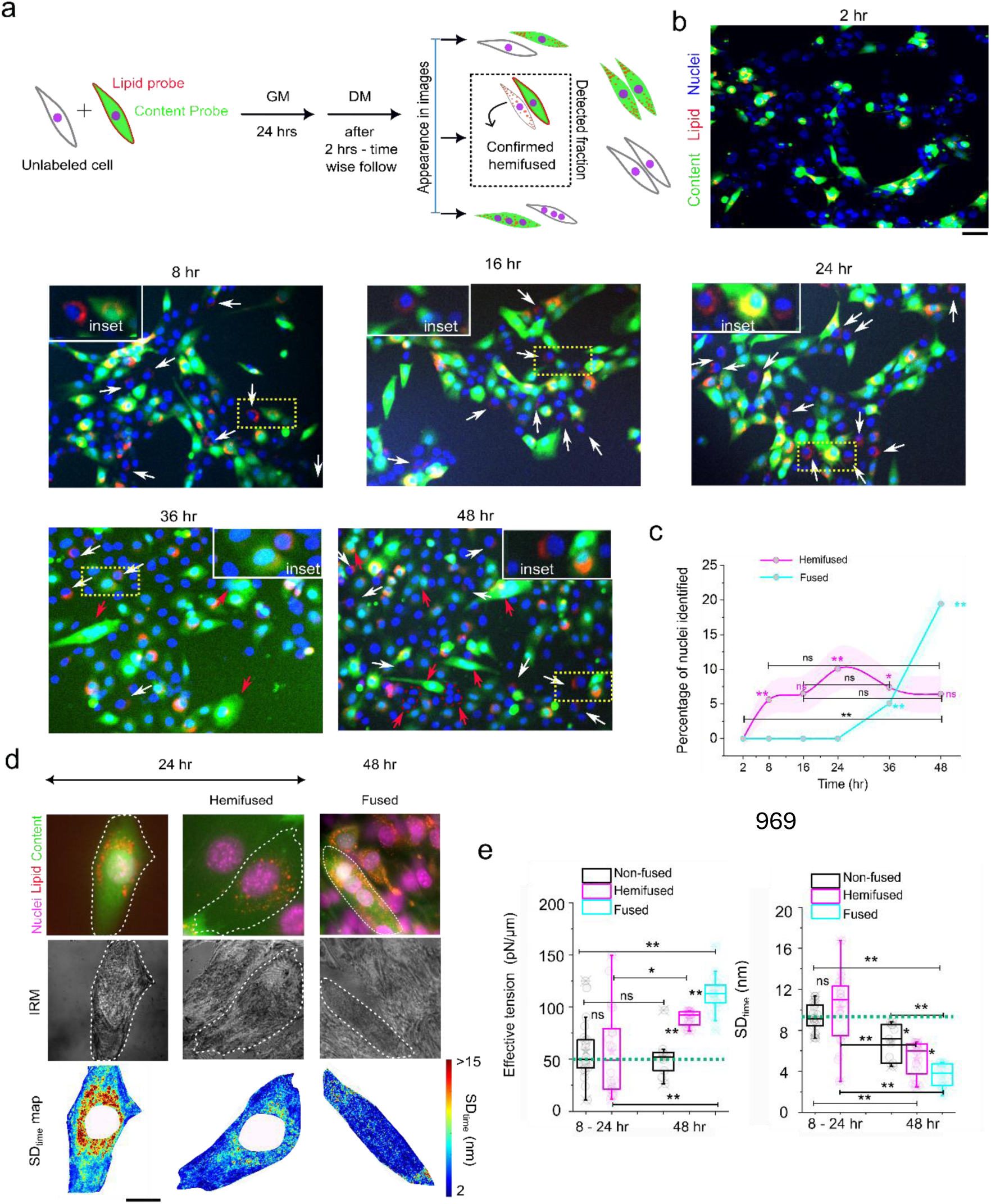
The early low-tension regime of myogenesis aligns with the hemi-fusing population. a) The schematic for differentially labelling the nonfused, hemifused and fused C2C12 cells. b) Representative (green – content probe, red - lipid probe, blue – (DAPI-stained) nuclei) epifluorescence images after 2 hr, 8hr, 16 hr, 24 hr, 36 hr and 48 hr of DM addition. White arrows mark the hemi-fused cells, and red arrows mark the fused cells. Insets represent typical hemifused cells. Scale bar – 10 µm c) Change in percentage of nuclei in hemifused and fused population of cells with tie after DM addition. Data are median ± s.e.m. (shaded region). N_field_ - 2 hr – 79, 8 hr – 68, 16 hr – 85, 24 hr – 120, 36 hr - 48,- 48 hr – 82. d) Representative images of sequential live Epifluorescence (top panel), IRM imaging (middle panel) and SD_time_ map of the membrane fluctuations of the respective cells (bottom panel). e) Effective tension and SD_time_ of nonfused, hemi-fused and fused cells after 24hr and 48 hr of DM administration. N_cell_ - 2 hr – 18 (non-fused), 25 (hemifused); 48 hrs- 11(non-fused), 26 (hemifused), 20 (fused) * - p<0.05, ** - p<0.001, ns - p>0.05. Mann-Whitney U-test performed, Scale bar – 10 µm. N_repeat_ −3.

Measuring percentage hemifusion with time - after knocking down Myomerger (**Fig. S2 a-c**) - did not show any difference from control at 24 hr, as expected. However, knocking down Myomaker suppressed hemifusion both at early and late timepoints confirming their individual roles. Although hemifusion events started at 8 and 24 hr, it was surprising that no fused cells were observed before 48 hr. To understand if Myomerger was active at 24 hr, the very early state of fusion was next identified by mixing cells either with red or green content-probe. Early fusion events when cell body or nuclei had no alignment were expected to be identified as “yellow” (leaking of red and green probes) cells (**Fig. S2 d, e**). Counting showed that while hemifusion started at 8 hr and peaked at 24 hr, fusion events only started at 24 hr and increased at/after 36 hr. Thus, despite Myomerger’s expression, its Myomerger’s ability to initiate fusion could be achieved at 24 hr and not earlier. At these timepoints, we next measured the state of tension in hemifusing cells.

IRM-based imaging provided mechanical information while fluorescence imaging identified the fusion-state of cells (**Fig. 1d**). IRM probes (**Fig. S1 a-e**) the nanometric height fluctuations of the basal membrane. After proper calibration (**Fig. S1 b-c**), membrane fluctuations amplitude was quantified as SD_time_ (**Fig. S1d)**. Effective membrane tension (and other parameters) was performed by fitting the power spectral density (PSD) (**Fig. S1e)** with Helfrich-based model (Biswas et al., 2017; Chakraborty et al., 2022). The fluctuation tension (termed “effective tension” in this study), thus derived, is different than the bare membrane tension or frame tension (Shiba et al., 2016). Nevertheless, recently, a significant correlation of IRM-based tension with apparent tension (measured by tether-extraction-based method) has been reported (Ghosh et al., 2026).

We observed that hemifused cells displayed similar tension as non-fused ones at 8/24 hr but progressively increased tension such that at 48 hr hemifused cells had higher tension/ lower fluctuations (**Fig. 1d, e**) relative to non-fused cells. The 2 - 24 hr phase was thus termed as the early phase of differentiation is distinct mechanically from 48 hr (**Fig. 1e**) for hemifusing cells. Multinucleated fused cells were observed only at 36 hr, 48 hr, and later time points. These cells had lesser membrane fluctuation amplitude and higher tension. These time points were therefore termed as the late phase of differentiation.

To understand if Myomerger had any role in increasing trend of global tension as differentiation progressed, knockdown studies were next performed.

### Myomerger switches from a membrane-relaxer to membrane-tensing agent with time

Knockdown of Myomerger by siRNA lowered fusion propensity of C2C12 cells by ∼60% (**Fig. 2a, S3a**); decreased its surface expression (**Fig. S3b**) and decreased the overall abundance of Myomerger (by ∼70% (**Fig. S3c**). We found that Myomerger knockdown increased the effective basal membrane tension by ∼25% (**Fig. 2a**, **Fig. 2b**) with a concomitant decrease in the amplitude of fluctuations (**Fig. 2c)** and increase in other parameters like confinement/effective viscosity/activity (**Fig. 2d-e, Fig. S3d**) for cells at 2/24 hr after DM treatment. However, at 48 – 72 hr, cells with Myomerger knocked down displayed lowered tension and higher membrane fluctuations (**Fig. S3e-g**). This late phase role of Myomerger was in line with previous reports which proposed that Myomerger helps in fusion pore formation post-hemi-fusion by increasing mechanical stress (Golani et al., 2021; Kozlov & Chernomordik, 2015). In contrast to Myomerger, siRNA Myomaker reduced overall cell tension and made the cell as well as the population more homogeneous for both the early and late phase (**Fig. S4**). Thus, the effect of Myomerger in the hemi-fusing stage is mechanically opposite to Myomaker’s effect on the overall cellular mechanics.

**Figure 2.**
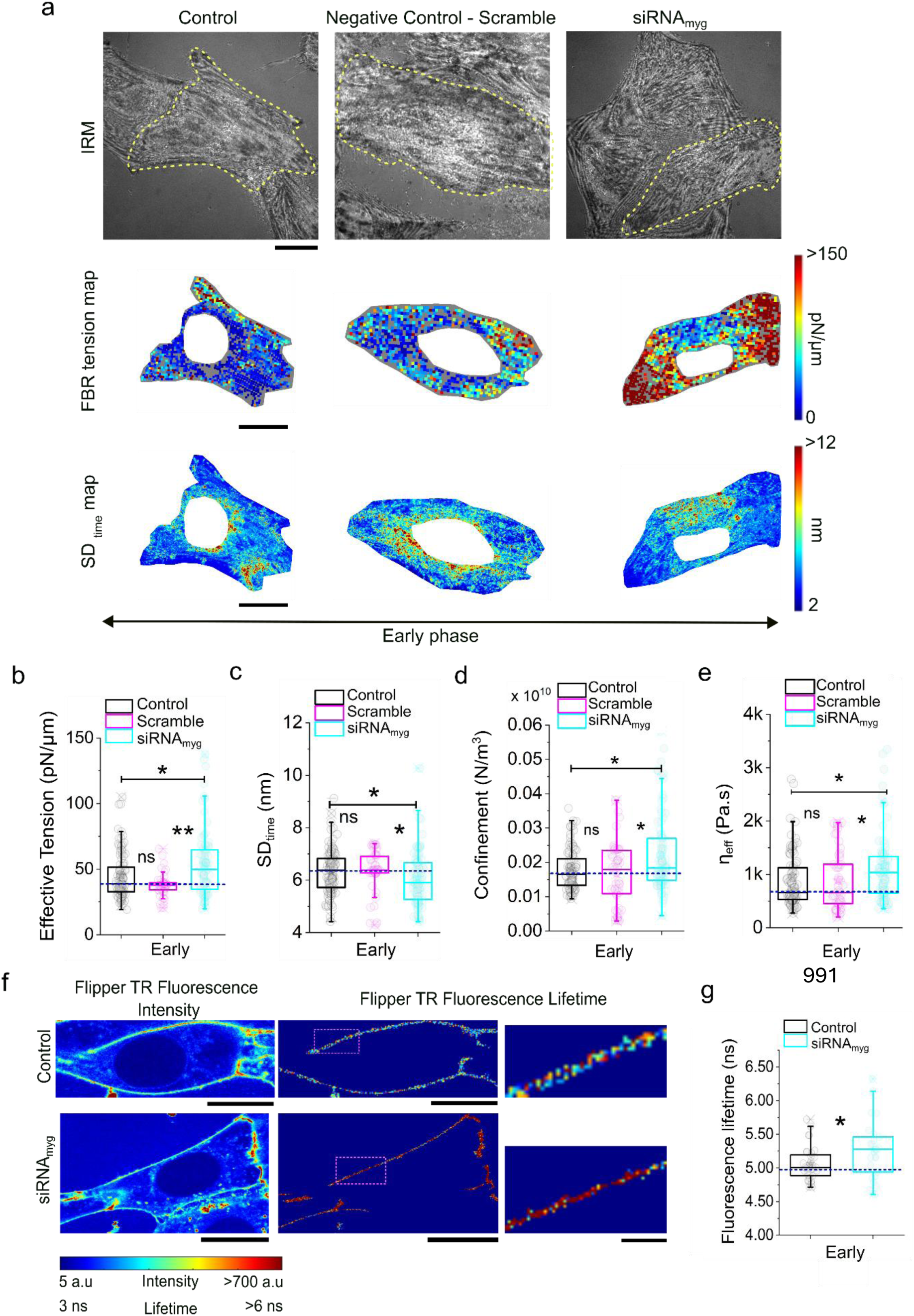
Myomerger aids in decreasing the basal membrane tension and lipid compaction globally in early phase. a) Left: Schematic of preparation for imaging; Right: Representative IRM images (top panel) and corresponding FBR-wise effective tension maps (bottom panel) of C2C12 myoblast cells in early phase (2 and 24 hrs post DM) of myogenesis, for control (untreated), negative control (scramble siRNA) and Myomerger knockdown (siRNA_myg_ treated) cells. (middle panel) b-e) Comparison of parameters measured from IRM imaging - effective tension, SD_time_, Confinement parameter and effective viscosity (*η_eff_*). N_cell_ – control (108), Negative control (Scramble) (57), siRNA Myomerger (cyan) (86) from 4 independent sets. Scale bar – 10 µm. f) Representative images of Flipper-TR Fluorescence Intensity and Flipper -TR Fluorescence lifetime. Scale bar – 10 µm. The insets marked in rectangular magenta boxes are zoomed in to depict the Flipper TR lifetime from the membrane region of the cells (within 1.5 µm above the basal membrane). Scale bar – 2 µm. g) Comparative study of Flipper -TR fluorescence lifetime of control versus siRNA_myg_ treated cells. The dotted blue lines mark the median of the control cells. N_cell_ – control (black) (32), siRNA Myomerger (cyan) (37). The error is SD in the box plots. * - p<0.05, ** - p<0.001, ns - p>0.05. Mann-Whitney U-test performed, N_repeat_ −3.

In order to further validate the observations on the cell surface mechanical state, Flipper-TR (**Fig. S3h**) lifetime (Colom et al., 2018) measurements were performed under similar conditions. Myomerger knockdown increased Flipper-TR’s lifetime significantly by ∼5% (**Fig. 2 f, g**). Under unaltered lipid microenvironment, lipid compaction and Flipper-TR lifetime are correlated to membrane mechanics (Colom et al., 2018). The decreased trend of lipid compaction in control cells than the Myomerger-depleted (knockdown) cells, could be due to the decreased tension (lower Flipper-TR fluorescence lifetime) of the PM (not only at the basal plane but also in midplanes) in the early time phase. However, this can be concluded only after the evaluation of the baseline lipid profile (e.g. cholesterol profiling).

These findings confirm that, during the early phase of differentiation, Myomerger contributes to reducing the effective fluctuation tension of cells, consistent with the observation that hemifusing cells exhibit a smaller tension difference than non-fused cells at this stage. Since early fusion events are known to commence around 24 h, we next sought to investigate the mechanism underlying Myomerger’s early function by examining its surface distribution and its interplay with membrane fluctuations.

### Super-resolution microscopy reveals Myomerger clusters at the basal PM

To understand the mechanism by which Myomerger regulates early membrane tension, the clustering pattern of endogenous Myomerger at the surface of the PM at the early phase was studied using immunofluorescence and super resolution STED microscopy. A 2D variant of STED was utilized with a lateral resolution of ∼ 30 nm. Cells were imaged at 2 and 24 hr post DM addition (**Fig. 3a, S5a, b**) and the fluorescent puncta observed were analyzed using object-detection algorithms. Clusters displayed majorly circular morphology except a small percentage of ring-like structures also observed (**Fig. S5c**). The distribution of the estimated equivalent diameter of the clusters displayed a bimodal distribution with peaks at ∼ 50 and 75 nm (**Fig. 3b**) indicating the basic structures to be of ∼ 40 nm diameter (estimating from a convolution-look-up plot, **Fig. 3b inset**) and another major population of doublets (∼ 70 nm). On plotting while the total intensities of the objects, presence of basic structures, doublets and higher order clusters with roughly double and higher intensities respectively were observed when checked in particular single cells (**Fig. 3c**). with an expected square-dependence of total intensity on diameter (**Fig. 3c inset**). Myomerger clusters has similar area footprint as Myomaker per unit area (area fraction) (**Fig. 3d**) and displayed a significant but very small percentage of overlap with Myomaker. It must be noted that we quantified the expected interference from non-specific contributions (**Fig. S5d**). A very low level 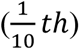 of maximum intensities was detected for background regions and samples with only secondary antibody staining (**Fig. S5d, e**) in comparison to real signals (**Fig. 3a**).

**Figure 3.**
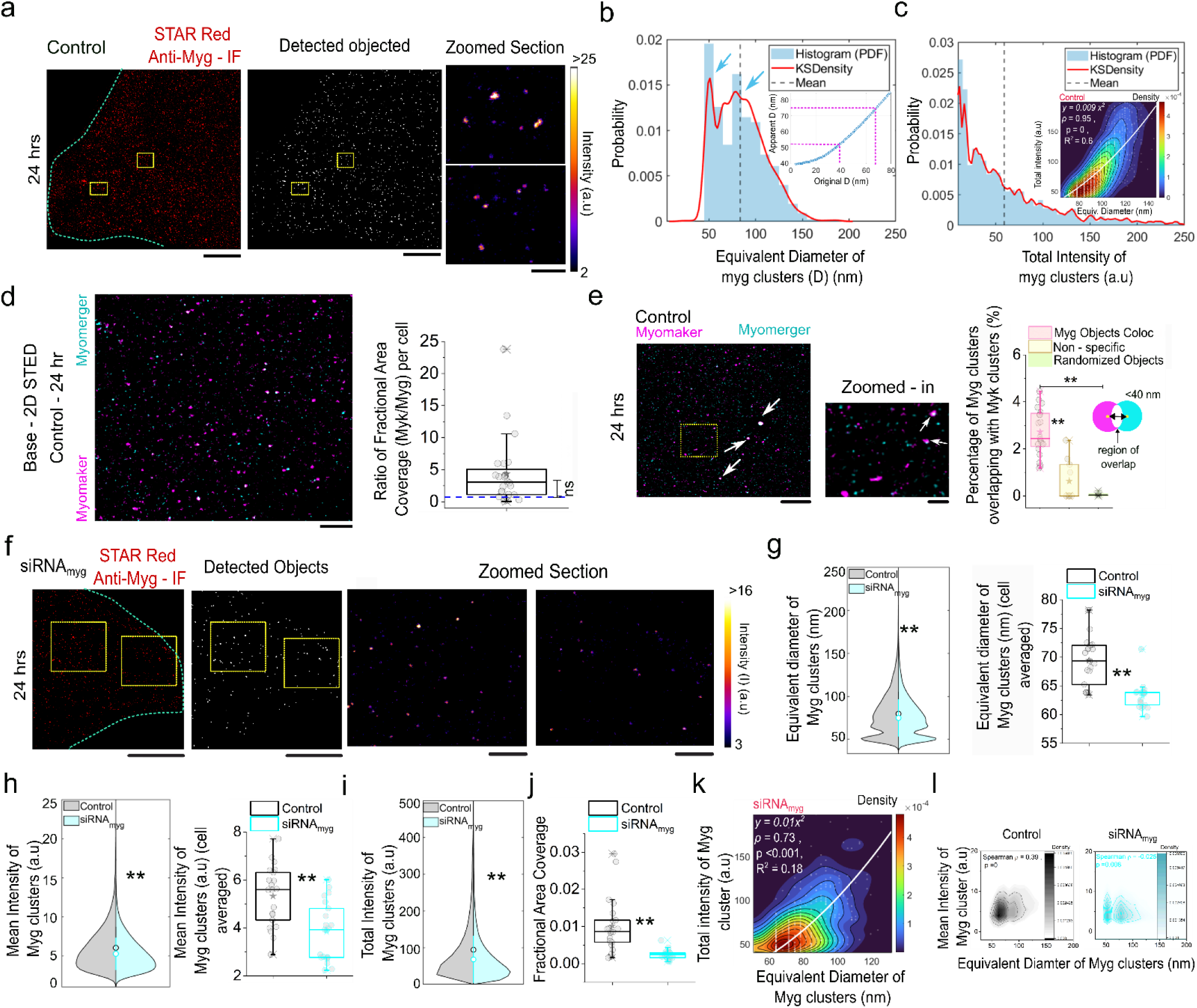
Super-resolution microscopy reveals Myomerger clusters at the basal plasma membrane of C2C12 cells at the early phase of myogenesis. a) Representative 2D STED images of endogenous Myomerger at basal surface of C2C12 cells using immunofluorescence. b) Normalized probability distribution (bandwidth – 10 nm) of object diameters with the corresponding kernel density estimate (red line); Cyan arrows point out the peaks; black dotted line depict the mean. Inset: convolution look up table with magenta lines drawn at observed (apparent) diameter to infer original diameter. c) normalized probability distribution of total intensities of Myomerger objects detection of a single representative cell. Inset: Total intensity vs. equivalent diameter; displays fit to y = ax^2^ fitted (white line) and Spearman correlation coefficient (ρ) and corresponding p value. N_repeat_ – 2, N_cell_ (control) – 21. d) Left panel: Representative 2D STED image of endogenous Myomerger and Myomaker. Right panel: Ratio of fractional area coverage of Myomaker clusters to Myomerger clusters. The significance test of median from 1 is noted. e) Representative image, Zoomed-in image and quantification of colocalization of Myomerger and Myomaker clusters at basal membrane using 2D STED. Yellow ROIs are regions zoomed in with white arrows showing colocalised clusters. Quantification involves counting clusters with centroid-to-centroid distances of less than 40 nm as overlapped (pink – actual Myg objects, yellow – non-specific background overlap, green – randomized Myg objects with Myk). N_repeat_ – 2, N_cell_ – 20. . f-j) Representative images and object-detection-based-quantification of effect of si-RNA-based Myomerger knockdown. N_cells_ : 21 control (black) and 21 siRNA_myg_ (cyan). N_clusters_ – control (18,206), siRNA_myg_ (10,856). (k) Depicts the y = ax^2^ fitted (white line) contour plot for total Intensities of Myomerger clusters (y axis) as a function of its equivalent diameter (x axis) under siRNA_myg_ treatment. l) 2D kernel density contour plots illustrate the control cells 24 hours post-DM treatment, and Myomerger knockdown conditions (black, cyan). The contours represent levels of estimated probability density obtained through kernel density estimation, displayed as a function of equivalent diameter (in nanometers) on the x-axis and mean intensity on the y-axis for Myomerger clusters. Regions with higher density indicate areas with a larger concentration of data points, as shown by the density scale bar. * p < 0.05, ** p < 0.001, ns p > 0.05. The Mann-Whitney U-test was used for analysis. For all box plot, error bar is SD. For all violin plots error bar is s.e.m. Scale bars : 5 µm, 500 nm (for zoomed-in section). Throughout the figure cyan dotted lines indicate cell boundaries; yellow doted lines mark out ROI used for Zoomed-in representations. Immunofluoresence used in this figure: Myomerger (Abberior STAR Red – cyan LUT) and Myomaker (Alexa Fluor 568 – magenta LUT).

Although the data clearly demonstrate Myomerger’s tendency to cluster but it also indicates tendency of clusters (or the basic structures) to aggregate rather than single clusters growing gradually bigger.

Reduction of Myomerger levels by siRNA treatment reduced the size, intensity and coverage of clusters (**Fig. 3f-j**). While total intensity retained the square-dependence on diameter, presence of doublets were clearer from mean intensity-diameter scatter plots. This part shows that at lower expression levels of Myomerger, reduces the size intensity – but clusters still can form doublets.

To understand the connection between the clustering ability of Myomerger with its role in membrane softening at the early phase, we next investigate the direct impact of these clusters on the underlying local membrane topology.

### Clustering results in membrane bending

To evaluate the local impact of Myomerger clustering on the basal plasma membrane of C2C12 cells, sequential correlative imaging approach combining TIRF-IRM (Chatterjee et al., 2025; Pattanayak et al., 2025) was adopted (**Fig. S6**). Live cells, transfected with fluorescently tagged Myomerger (tGFP-Myg) were imaged via TIRF – IRM at 2 and 24 hr (early phase) post DM administration. TIRF enabled imaging of tGFP-Myomerger clusters expressed on the basal membrane – due to low penetration depth of ∼ 100 nm of the evanescent-mode-while sequential IRM at the same region enabled mapping of the amplitude of membrane fluctuation (SD_time_) (**Fig. 4a-b**). From the TIRF time series of a single cell captured at 3.33 frames per sec for 300 frames, the membrane-localized, immobile Myomerger clusters were identified using kymograph (**Fig. 4c**) of TIRF intensity in contrast to any mobile vesicular form.

**Figure 4.**
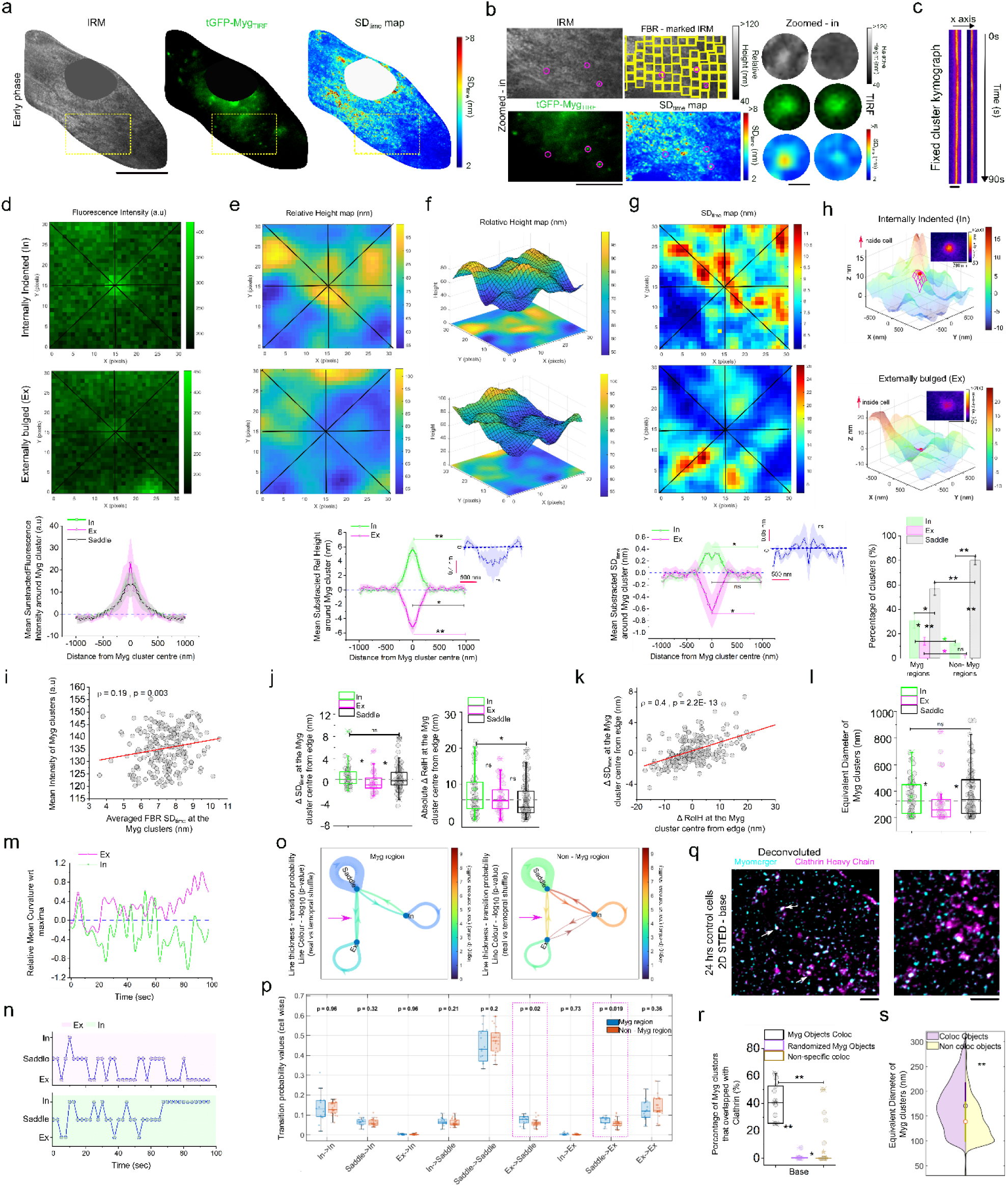
Membrane bending topology correlates with Myomerger clustering in a size/density dependent manner. a) Representative images of early phase tGFP-Myomerger transfected C2C12 cells were imaged using sequential correlative TIRF-IRM. The left panel shows a typical IRM basal membrane image, with the middle panel showing Myomerger expression. The right panel shows a pixel-wise map of membrane height fluctuations (SD_time_). The nucleus is excluded, and the yellow dotted rectangle indicates a zoomed-in inset. Scale bar – 10 µm. (b) IRM relative height map of ROI marked out in (a) and zoomed in images of relative height, fluorescence and SD_time_ map with magenta circles marking fixed clusters used for topology mapping. Scale bar – 5 µm. Right Scale bar – 600 nm. (c) Kymograph for typical fixed clusters from 300 ms movies of 300 frames. (d-g top and middle panels) d)Display fluorescence intensity (e,f), relative height (g) SD_time_ map and (bottom panels) radially averaged profiles around Myomerger cluster centers for different membrane topologies (In – green, Ex – magenta, Flat – black). (h) (top panel) Surface plot of a 31×31 membrane patch around a Myomerger cluster (red blob), showing curvature fitted (second order polynomial) (magenta mesh) to a 5×5 patch, indicating inward (top) or outward (bottom) bulges, with the red arrow indicating the cytoplasmic side. (bottom panel); (bottom panel) Percentage of clusters displaying In, Ex, and Flat topologies obtained from curvature amps at cluster centres compared with Non-Myg regions from the same cells (non-fluorescent regions). (i) dependence of mean intensity of Myomerger clusters on FBR-wise averaged SD_time_. The Spearman correlation coefficient (ρ) and p value is noted on the plot. The red bold line represents the linear fit. (j) (left panel)) Comparison of ΔSD_time_ plot at cluster centres versus edges (within 1 µm). (right panel) Absolute Δ Rel Height at cluster centres versus edges (within 1 µm). (k) Dependence of ΔSD_time_ at clusters centre on corresponding ΔRel Height. The Spearman correlation coefficient (ρ) and p value is noted on the plot. (l) Comparison of Equivalent Diameter of Myomerger clusters at In, Ex, and saddle features of the membrane. (m) Time series of Relative curvature normalized wrt maximum curvature generated at Myomerger clusters centres at In (green) and Ex (magenta) features of the membrane (quantification of over every 50 frames (2 secs)). (n) Time series of class detected (o) (left panel) transition probability network for myomerger-enriched region and non-myomerger regions. The thickness of lines depicts the strength of the probability, and the colour bar represents the p value for significance with the randomised transition states (the sequence of consecutive transition states was reshuffled in random). The magenta arrow shows the transitions that are significant in real Myomerger cluster regions wrt to the Non-Myomerger cluster regions (p) Comparison of transitions with jitter points denoting averaged transition probability values per cell at Myomerger cluster vs Non cluster regions. (q) The Myomerger (cyan – IF-Abberior STAR red) and clathrin heavy chain (magenta – IF – Alexa Fluor 568) clusters co-distribution at the basal membrane of 24 hr DM treated C2C12 cells captured by 2D STED. The white arrows denote the clusters having colocalization. right: zoomed-in representation. Scale bar – 5,3 µm. (r) Comparison of overlap (s) Comparison of equivalent diameters. N_repeat_ −2, N_cell_ – 21, N_cluster_ - 4642, * p < 0.05, ** p < 0.001, ns p > 0.05. The Mann-Whitney U-test was used for analysis. For all line plots, the median line profile with s.e.m as the shaded region is depicted. For all box plots, error is SD.

To estimate the local impact of Myomerger cluster on the membrane topology, fluorescence cluster centres are detected by object-detection algorithms and the analysis subsequently focused around the centres (**Fig. 4d**). Linescans (**black lines, Fig. 4d**) around the cluster centers are performed for fluorescence as well as IRM images. From IRM images, we obtained the maps of mean relative height (averaging over all timeframes) (**Fig. 4e, f**) and SD_time_ (**Fig. 4g**). Linescans – radially averaged (see methods) - yield the profile of each quantity. At every fluorescence peak (**Fig. 4d bottom**) the corresponding membrane topology (**Fig. 4e,f**) and fluctuations (**Fig. 4g**) were studied. The topology of membrane (5×5 pixels) (**Fig. 4h, red wire frame**) around each cluster centre were analyzed by extracting the mean and Gaussian curvatures around the cluster centres (from mean relative height from IRM images) and classifying them as internally-indented (In) /externally-bulged (Ex) and Saddles using those parameters (see Methods) (**Fig. S7(a-b)**). The averaged profiles directly plotted confirm the In/Ex relative height to be as expected (**Fig. 4e bottom**) while when averaged across clusters revealed no specific curvature (**Fig. 4e bottom inset**). While highest represented class was saddles (**Fig. 4h bottom**), they were lesser than found at other regions not specifically enriched in Myomerger. Saddle regions when radially averaged, revealed a shallow external bulge (**Fig. S7c**). In and Ex classes were present more than for other regions. However, among them although Ex class was sparser than In class for Myomerger clusters, they were very rare in non-myomerger regions (**Fig. 4h bottom**). Membrane fluctuations were damped for Ex clusters while enhanced at In clusters and was unaffected at Saddles (**Fig. 4g bottom, S7d**). The level of damping or enhanced fluctuations induced by Myomerger clusters were not significantly different from the non-myomerger regions, though by itself these events were rare in non-myomerger regions (**Fig. S7e-f**). Overall, clusters having higher intensity also displayed higher fluctuations at their centre, suggesting In and Ex may have different sizes. While fluctuations varied (**Fig. 4j left**), the depth of the indentations (plotted as change in relative height: ΔRelH – **Fig. 4j right**) didn’t show significant differences between classes. However, ignoring classification, dampening/enhancing of fluctuations, the centre was clearly correlated with the depth at the centre (**Fig. 4k**). The equivalent diameters were found to be higher for In-class or Saddle-class clusters than the Ex-class (**Fig. 4l**). It was also observed that the classes had similar goodness of fit irrespective of their curvature state and a threshold of 0.7 as goodness of fit was considered for analysis (**Fig. S7g**). We also studied that if classified through IRM imaging, if clusters transitioned from one class to another. While typical time-series (**Fig. 4m**) showed that once previously classified as In / Ex mostly remained of similarly curved most part of the 108 secs imaged, we next classified for every 2 seconds (**Fig. 4n**) and quantified the transitions between classes (**Fig. 4 o, p**). Each class showed highest probability to remain the same class (**Fig. 4 o – loops**); however, each class had similar stability as found non-myomerger region. Most transitions were allowed except that between In and Ex – in contrast to the transition quantified for non-myomerger regions. Saddle-Ex transitions were also significantly enhanced than in non-myomerger regions (**Fig. 4 o – magenta arrow, Fig. 4p**).

To understand the significance of the In-class structures, we evaluated the colocalization of Myomerger clusters with Clathrin heavy chain (CHC) (**Fig. 4q, S7h**). We observed ∼ 40% Myomerger clusters to have overlaps with CHC clusters (**Fig. 4r**). Interestingly the observed percentage of In from TIRF-IRM studies were also ∼ 32% (**Fig. 4h bottom**). Myomerger clusters that colocalized with CHC clusters also displayed a significantly bigger size as In-class structures displayed in comparison (**Fig. 4l**).

Thus, we establish the regions with Myomerger enrichment get stable curvatures but while at smaller sizes they ex-vaginate as size grows they go from saddle/flat structures to invaginations – probably to get endocytosed by Clathrin mediated pathway. To validate similar membrane remodelling in endogenous Myomerger clusters, we next study the local remodelling of lipids by evaluating Flipper-TR’s lifetime.

### Local lipid compaction profiles at Myomerger clusters reveal clustering stabilizes curvatures at the neck of clustering

To further probe the local lipid compaction induced by the endogenous Myomerger molecules simultaneous STED-FLIM using Flipper-TR probe and living cell antibody staining was performed. STED-FLIM was performed at both the basal planes and at midplanes to characterize clustering with lipid compaction at both the basal plane and edge membranes at midplanes. (**Fig. 5a**). To minimize photo-toxicity bigger pixel-size (thus faster imaging) was used and the detected cluster diameters reveal very small cluster, one population with peak ∼ 300 nm and tail after 400 nm (**Fig. S8(a,b)**. Through radially averaged profiles we show that, for smaller clusters lesser than 400 nm diameter (similar to smaller Ex-class clusters), lifetime is higher towards the centre with a lowering at the edge (**Fig. 5b**). Individual clusters show similar peaking profile (**Fig. 5c**). For bigger clusters, a lowering of lifetime at the centre was noted (**Fig. S8c**). Quantifying the intensity-lifetime correlation at each cluster, while a positive Spearman correlation coefficient was observed for smaller clusters, bigger ones displayed a lower coefficient (**Fig. S8b**). Even at the lateral membrane at an equatorial plane (not cell base), Flipper-TR probe at Myomerger clusters reveal significant higher lifetime at the centre and reduced compaction at the cluster-edge (**Fig. 5e**). No size-based pooling was done because the average size of clusters here were bigger than at the base.

**Figure 5.**
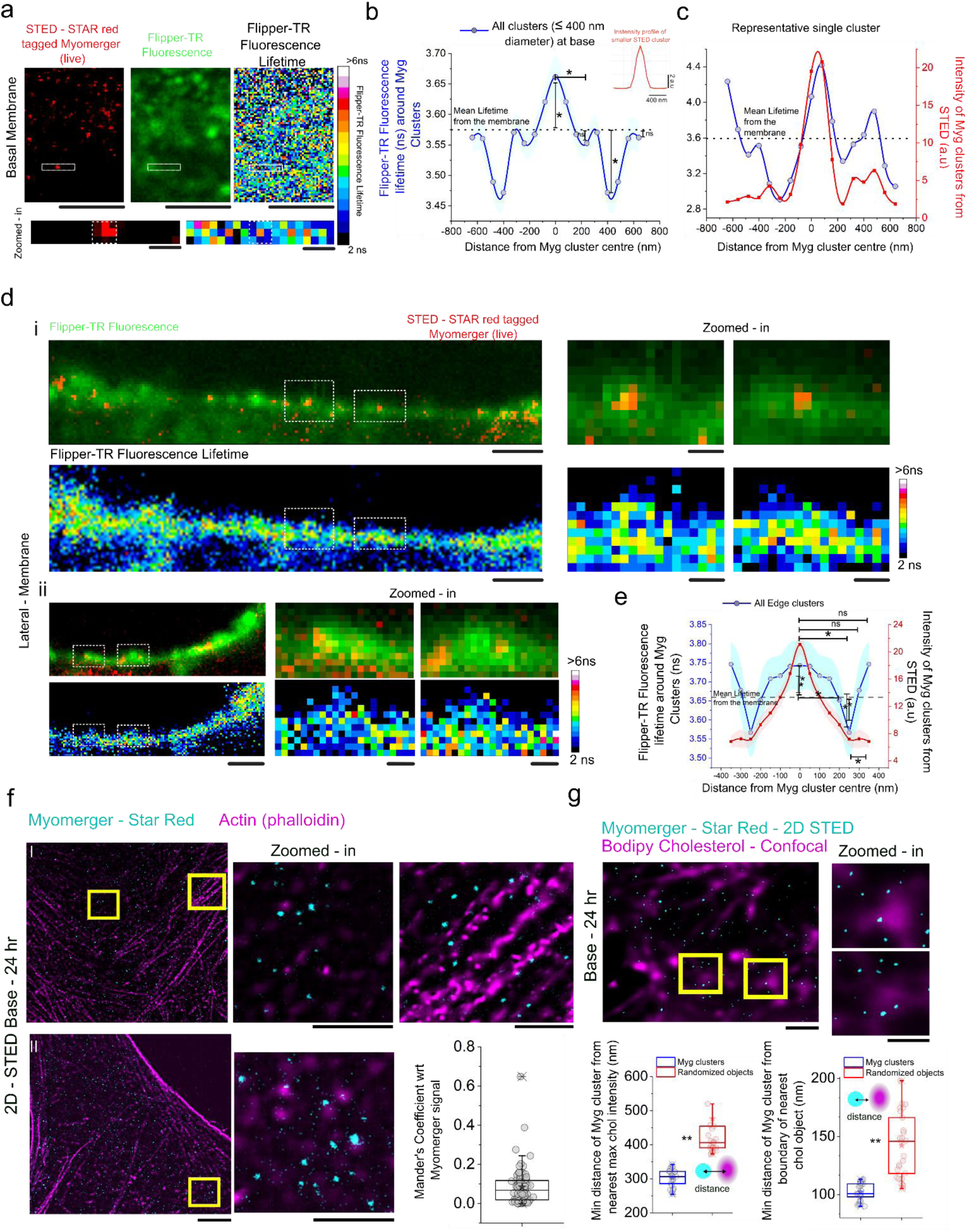
Local lipid compaction profiles at Myomerger clusters reveal clustering stabilises curvatures at the neck of clustering. a) Representative 2D live STED C2C12 early phase basal membrane section (left) with corresponding simultaneous FlipperTR-fluorescence intensity (middle) and lifetime (right). Scale bar – 2 µm. The inset marks a cluster of Myomerger that is zoomed in the bottom panel to observe the local Flipper-TR lifetime trend at Myomerger clusters due to lipid compaction. Scale bar – 200 nm. The white dotted box denotes the cluster position. (b) The symmetric profile of Flipper-TR lifetime from the basal membrane centred around the smaller Myomerger cluster (< 400 nm in diameter). The inset shows the average radial profile of fluorescence intensity of Myomerger clusters. 0 in x-axis marks the centroid of cluster. Dotted black line marks the mean lifetime of the basal membrane. (c) A representative raw double-y plot to represent the Fluorescence intensity of Myomerger cluster (red) and its corresponding region Flipper-TR Fluorescence Lifetime around the cluster centre(blue). ((d) (i) and (ii) (top panel) Merged Representative images that depicts the lateral membrane of early phase C2C12 cells (at z sections above base). Green channel represents the Flipper TR Fluorescence and Red channel represents the STAR red - immunostained live cells against endogenous Myomerger clusters. (bottom panel) The corresponding Flipper-TR lifetime map. The insets are marked by white square rois. Scale bar – 1 µm. The marked rois are zoomed in. Scale bar – 200 nm. Top panel shows the position of Myomerger cluster wrt to edge membrane, along with Flipper-TR probe. Bottom panel denotes the Flipper-TR Fluorescence lifetime. (e) The Double y symmetric profile of Flipper-TR lifetime from the basal membrane centred around the Myomerger cluster that are at the lateral membrane. 0 in x axis marks the centroid of Myomerger cluster. Dotted black line marks the mean lifetime of the basal membrane. N_repeat_ −2, N_cell_ – 10, N_cluster_ – 258 (basal membrane) (smaller clusters– 178); 79 (edge of basal membrane). (f) (I)(II) Representative section from 2D STED imaging of the C2C12 cell basal membrane, taken 24 hours after DM administration. Fixed cells are immunolabeled for endogenous Myomerger (Abberior STAR Red – cyan LUT) and F-actin (Alexa Fluor 568 Phalloidin – magenta LUT). The yellow rectangular insets are zoomed in. The Manders coefficient of the Colocalization of Myomerger clusters with F-actin wrt total Myomerger signal from cell is depicted. Scale bar 5, µm 500 nm. (g)(top panel) Representative section from 2D STED - confocal imaging of the C2C12 cell basal membrane, taken 24 hours after DM administration. Fixed cells are immunolabeled for endogenous Myomerger (Abberior STAR Red - 2D STED – cyan LUT) and cholesterol (Bodipy cholesterol – confocal -magenta LUT). The yellow rectangular insets are zoomed in. Scale bar 1 µm, 500 nm. (bottom panel) (left) The comparative study of minimum distance of real Myomerger cluster centre from highest Intensity peak from the nearest cholesterol object versus the randomised Myomerger cluster centroid positions. (right) The comparative study of minimum distance of real Myomerger cluster centre from boundary of the nearest cholesterol object versus the randomised Myomerger cluster centroid positions. N_repeat_ −2, N_cell_ – 23 (F-actin-Myg coloc); 25 (chol - Myg coloc), The Mann-Whitney U-test was used for analysis. * p < 0.05, ** p < 0.001, ns p > 0.05. For all line plots, the median line profile with s.e.m as the shaded region is depicted. For all box plots, error is SD.

Myomerger is known to induce positive curvature on liposome membranes (Golani et al., 2021). In this section we show that smaller clusters indeed induce an externally bulged topology on cell membrane while bringing in sharp changes lipid compaction in its vicinity both at basal and lateral membranes.

### Myomerger clusters remain only close to actin/cholesterol enriched regions

Having demonstrated membrane remodelling at Myomerger clusters, we next probed if such remodelling could be explained by direct action of actin/cholesterol enrichment at Myomerger clusters. Labelling F-actin with Phalloidin, STED imaging was performed at basal cell membranes with cortical actin captured together with IF of endogenous Myomerger (**Fig. 5f, Fig. S8d**). No visible overlap could be seen. On quantification, Mander’s coefficient was found to indicate the same (**Fig. 5f**). However, on calculating the minimum distance of cluster centre from the boundary pixels of actin objects and comparing with a similar calculation but using randomized positions of Myomerger objects, we observed a clear reduction in the minimum distance. Hence, although actin did not localize at cluster centres pushing/pulling them, clusters tended to localize next to actin filaments. Cholesterol (labelled with Bodipy-Chol and imaged in confocal) on the basal membrane was observed to have microscopic domains of enrichment and Myomerger remained at their edges and not colocalizing at the centres (with maximum intensity). However, again, Myomerger clusters were closer to the centres/boundary pixels of Chol-objects than expected from a random distribution of Myomerger (**Fig. 5g, S8e**).

Together we show that membrane bending or lipid compaction by Myomerger need not be through recruitment of actin/cholesterol.

## Discussion

Cell-cell fusion is at the heart of skeletal muscle formation and repair and defining how muscle fusogens are regulated is a prerequisite for manipulating fusion therapeutically. However, previous literature primarily talks about the predominant role of Myomerger as a membrane stressor during the hemi-fusion to permit fusion pore formation process (Chen et al., 2020; Golani et al., 2021; Kozlov & Chernomordik, 2015). By resolving hemifusion and content mixing as separate events, and without imposing any biochemical constraint on the system - no LysoPC or other fusion-arresting agent - we find that hemifusion begins early and accumulates slowly, reaching ∼5% of cells by 8 hr and ∼10% by 24 hr, whereas the earliest content-mixing events appear only at 24 hr and multinucleated cells only from 36 hr. We show that Myomerger is already expressed at 2 hr/8 hr yet cannot convert a hemifusion intermediate into a pore until roughly 24 hr.

While Myomerger’s surface expression and regional tension had been shown to be negatively correlated at early phase of myogenesis (Chakraborty et al., 2022), we demonstrate causation though knockdown studies and reveal that Myomerger’s role evolves. Towards this, IRM was used as a non-invasive tool for the comparative study of cell surface mechanics. Because IRM is limited to the basal membrane, which has no active role in fusion for myogenesis, it helped in probing the global mechanical modulations induced by Myomerger. Our study on basal cell surface mechanics is therefore simultaneously complemented by an indirect Flipper-TR- lipid compaction dependent tension marker, by FLIM imaging. Flipper-TR fluorescence lifetime is known to be inversely related to membrane tension, in single phase L_o_ GUVs but in cellular systems and mixed-phase GUVs lifetime and tension are positively correlated(Colom et al., 2018). IRM measures effective cell surface tension that is influenced by the underlying cytoskeleton while Flipper-TR-based tension measurements considers only the membrane tension which might be affected by cholesterol like lipid content (Ragaller et al., 2024; Roffay et al., 2023). Therefore measurements at the mid z-planes (above the base) using Flipper-TR probe were used to complement IRM. So, agreement between the two readouts is best treated as corroboration by an orthogonal observable rather than as validation of either in the other’s.

Contrary to what a purely local model of hemifusion would predict, hemifusing cells were mechanically indistinguishable from their non-fused neighbours at 8 and 24 hr, and only became significantly more taut, with correspondingly damped fluctuations, by 48 hr (**Fig. 1d, e**). The mechanical signature of hemifusion is therefore not present when hemifusion is at its most frequent; it emerges later, as the population transitions to myotube formation. This separates differentiation into an early phase (2 - 24 hr) and a mechanically distinct late phase (36 hr onward), and it means the low tension of early-phase cells cannot be a consequence of hemifusion. We suggest instead that it is a permissive state that precedes it. Multinucleated cells, observed from 36 hr, carried the lowest fluctuation amplitudes and the highest tension of any population - recovering in mammalian myoblasts an effect described in Drosophila, where the receiving founder cell stiffens its actomyosin cortex (Kim et al., 2015).

The knockdown experiments subsequently establish that it is Myomerger (**Fig. 2, S3**) (not Myomaker (**Fig. S4**)) whose biphasic mechanical impact on the basal membrane is in line with the progressive build-up of membrane stress in hemifused cells post DM addition. However, the cell-level effects contrast the local impact measured and presented in (Fig. 4). Our attempt to probe the local mechanics at Myomerger-enriched regions depicts that Myomerger induces topological changes in a size-dependent manner (**Fig. 4**).

Averaged across all clusters, the membrane at Myomerger sites showed no net curvature (**Fig. 4e inset**). Myomerger may not impose a single deformation. What it might change is the distribution of curvature states. Classifying the local topology by mean and Gaussian curvature, saddles remained the commonest class at Myomerger sites but were less frequent there than at non-enriched regions, while both internally indented (In, ∼32%) and externally bulged (Ex) classes were over-represented. The Ex class is the more informative of the two: although sparser than In even at Myomerger clusters, it is very rare in non-enriched membrane, making outward bulging the topological signature specific to Myomerger. Ex clusters had smaller equivalent diameters and damped fluctuations, whereas In and saddle clusters were larger, with In clusters showing enhanced fluctuations. Across all clusters, fluctuation amplitude at the centre tracked the depth of the local deformation. It should be stressed that the magnitude of damping or enhancement at Myomerger clusters was not significantly different from the equivalent rare events at non-enriched regions (**Fig. S7e–f**). Myomerger makes these curvature states common rather than making them more extreme. However, making them more or less common would alter the overall mechanics.

The dynamics point the same way. Each class was most likely to persist, with a stability comparable to non-enriched regions, but the transition structure differed: In and Ex never interconverted directly at Myomerger clusters, whereas non-enriched regions permit that transition, and saddle-to-Ex transitions were significantly enhanced (**Fig. 4o, p**). Bulges and indentations at Myomerger sites are therefore not two ends of one continuum but separate states reached through a flatter intermediate – indicative of differing biological functions. Combined with the size dependence, this suggests a progression in which a small cluster bulges outward, and as it grows passes through a saddle state into an indentation. Consistent with that reading, ∼40% of Myomerger clusters overlapped clathrin heavy chain — close to the ∼32% In fraction - and clathrin-positive clusters were significantly larger, matching the In-class size. Endocytic removal of the largest clusters would cap how much Myomerger any one site can accumulate, and offers a concrete mechanism for the surface regulation of Myomerger invoked previously (Golani et al., 2021).

STED-FLIM of Flipper-TR with live antibody labelling of endogenous Myomerger provided an independent readout of the same size dependence. Clusters below ∼400 nm showed higher lifetime at the centre falling towards the edge (**Fig. 5b, c**), the opposite of the profile seen at larger clusters (**Fig. S8c**), and the intensity–lifetime correlation was positive for small clusters and weaker for large ones. The same central compaction was found at the lateral membrane in equatorial planes (**Fig. 5e**), so this is not an artefact of substrate proximity. The two assays converge: small clusters are locally compacted and locally damped, large clusters are neither. We note that the clusters detected under STED-FLIM are considerably larger than those measured by STED alone, a consequence of the coarser sampling used to limit phototoxicity, so the correspondence between the size classes in the two experiments is by analogy rather than by direct measurement.

Finally, we asked whether this remodelling could simply reflect recruitment of actin or cholesterol. Myomerger showed no visible overlap with phalloidin-labelled cortical actin and low Mander’s coefficients, yet cluster centres lay significantly closer to actin object boundaries than randomised cluster positions. The same held for cholesterol: Myomerger sat at the edges of Bodipy-cholesterol-enriched domains rather than at their intensity maxima, but closer to them than chance allows. It is therefore possible that the clusters occupy sites of pre-existing curvature, consistent with a curvature preference on Myomerger’s part, while arguing against actin or cholesterol enrichment being the agent of the bending we observe.

Taken together, these results establish that Myomerger assembles into discrete, quantised clusters at the plasma membrane during myogenesis, and that cluster size tracks local mechanical output: small clusters bulge outward and compact the local lipid environment, large clusters indent and are handed to the clathrin machinery. Cell-scale mechanics should therefore follow the evolving area fraction and size distribution of these clusters – though not of Myomerger’s alone, since Myomaker also modulated cellular mechanics in both phases. Since the current measurements remain correlative, disrupting or enhancing clustering and finding the shift in the time evolution of fusion-pore-formation would be critical to finally establish the causal link. We characterised clusters on the basal membrane of the general population, leaving their localisation and activity at hemifusion interfaces still to be established. The contribution, therefore, is the first demonstration of the effect of Myomerger clustering on cell membrane, and a new pair of parameters - cluster size and local membrane topology - through which Myomerger-dependent fusion can be dissected.

## Methodology

### Resources

| REAGENT or RESOURCE | SOURCE | IDENTIFIER |
| --- | --- | --- |
| **Antibodies** |  |  |
| Myosin 4 Monoclonal Antibody (mouse) (MF20) | eBioscience | Cat# 14-650-82; LOT# 4301341 |
| ESGP Polyclonal Antibody (against mouse raised in sheep) | Invitrogen | Cat# PA5-47639; LOT# UJ2868751A |
| Goat anti-Mouse IgG H&L, Alexa Fluor 488 | Abcam | Cat# ab150117; LOT# GR3192504-1 |
| Donkey anti-Sheep IgG (H+L) Secondary Antibody, Alexa Fluor 488 | Invitrogen | Cat# A11015; LOT# 2079356 |
| Goat anti-Rabbit IgG H&L, Alexa Fluor 568 | Abcam | Cat# ab175471 |
| Clathrin heavy chain Rabbit mAb | Abclonal | CAT#A4943; LOT#4000001228 |
| Alexa Fluor 568 Phalloidin | Life Technologies | CAT#A12380; LOT#1600223 |
| Anti-TMEM8C antibody (raised in Rabbit) | Abcam | Cat# ab188300 |
| Abberior STAR RED, donkey anti-sheep IgG, 500 µl (1 mg/ml) | Abberior | Cat# STRED-1056 |
| **Pharmacological reagents** |  |  |
| Flipper-TR Bioprobe | Spirochrome | CAT#CY-SC020; LOT#057 |
| Mowoil 4-88 | ALDRICH Chemistry | CAT#81381;<br>LOT#BCBK2317V |
| Tris Hydrochloride | SRL | CAT#99438; BATCH#<br>8677879 |
| DABCO 33-LV | ALDRICH Chemistry | CAT#290734;<br>LOT#MKBK8680V |
| Vybrant Dil (lipid probe) | Thermo-Fisher Scientific | CAT#V22885 |
| Green CMFDA cell tracker | Thermo-Fisher Scientific | CAT#C7025 |
| Orange CMRA cell tracker | Thermo-Fisher Scientific | CAT#C34551 |
| Bodipy cholesterol | MedChemExpress | CAT# HY-125746 |
| Gm7325 (Myomerger)<br>(tGFP-tagged Mymx-Mouse predicted gene 7325<br>(Gm7325) transcript variant<br>3 | OriGene | CAT#MG224234 |
| Magnesium Chloride<br>hexahydrate Cryst.<br>Purified | Merck | Cat# 442615; CAS# 7791-18-6; |
| HEPES | Sigma-Aldrich | Cat# H3375;<br>LOT#SLBN0452V |
| Sodium chloride | Sigma-Aldrich | Cat# S7653; LOT#<br>081M0051V |
| Dimethyl sulfoxide | Sigma-Aldrich | Cat# 276855; LOT#<br>SHBC3339V; |
| Lipofectamine™ 3000<br>Transfection Reagent | Invitrogen™ | CAT#L3000015 |
| Oleic acid | Sigma-Aldrich | CAT#O1008 |
| Myomerger siRNA<br>(mm.Ri.Gm7325.13.1)<br>(duplex pre-<br>designed siRNA) | IDT - 10 nmol | CAT#IDT27555 |
| **Cell culture and IF chemicals** |  |  |
| Dulbecco's Modified Eagle's<br>Medium (DMEM, High Glucose) | Gibco | Cat# 11965092 |
| Fetal Bovine Serum, certified, heat inactivated, US | Gibco | Cat# 10082147 |
| Antibiotic-Antimycotic (100X) | Gibco | Cat# 15240062 |
| Horse Serum, New Zealand origin | Gibco | Cat# 16050122 |
| Insulin solution human | Sigma-Aldrich | Cat# I9278; LOT# SLBX0973 |
| Trypsin-EDTA (0.25%), phenol red | Gibco | Cat# 25200072 |
| Paraformaldehyde | Sigma-Aldrich | Cat# P6148 |
| Phosphate Buffered Saline | Sigma-Aldrich | Cat# P3813 |
| Opti-MEM™ I Reduced Serum Medium | Gibco | Cat# 31985070 |
| Gelatin | Sigma-Aldrich | Cat# G2500; LOT# SLBX2973 |
| Glycine | Merck | Cat# 56-40-6 |
| TritonX-100 | Sigma-Aldrich | Cat# 101371902 |
| DAPI | Sigma-Aldrich | Cat# D9542; Batch# 0000081272 |
| **For calibration** |  |  |
| NIST Traceable Particle Size Standard, 60 µm | Bangs laboratories | Cat# L130806L; LOT# 11247 |
| **Software and algorithms** |  |  |
| ImageJ (FIJI) | NIH | https://fiji.sc/ |
| Origin | OriginLab Corporation | NA |
| MATLAB | The Mathworks, Inc. | 40684590 |

### Cell line

The mouse skeletal myoblast cell line (C2C12) from ATCC (CRL-1772) was used as a model for studying fusion and differentiation and the corresponding underlying mechanisms. Cells were checked for *Mycoplasma* and tested negative. Institutional Biosafety Committee approved the use of C2C12 cells.

### Cell culture

C2C12 cells were grown in growth media (GM). GM contained Dulbecco’s modified Eagle’s Medium (DMEM, Gibco, Life Technologies, United States) supplemented with 10% fetal bovine serum (FBS, Gibco, HI, US origin), and 1% antibiotic–antimycotic solution (Gibco). For induction of differentiation/myogenesis, the seeding density of C2C12 cells was maintained nearly at ∼60,000 cells/ml. Seeding was carried out in customized round glass-bottom dishes and maintained in GM until 70 - 80% confluency is attained. Following that GM was replaced with differentiation media (DM) consisting of DMEM, 2% horse serum (HS) (Gibco), 1% antibiotic–antimycotic solution, and 0.1% insulin (1 μM) (Sigma-Aldrich) solution. Cells were maintained at 37°C in a humidified incubator with 5% CO_2_ for 5 days in DM, and the medium was replaced every 24 h for obtaining myotubes as a result of differentiation. All experiments were performed with cells having passage number from 3 - 10. Beyond passage 10, differentiation into myotubes becomes difficult.

### Hemi-fusion and Fusion Pore Detection Assay

For estimating percentage of hemifusing cells, two populations of C2C12 cells of 75% confluence were first grown separately. Subsequently, one population was incubated with only DMEM with Hoechst dye, and the other population was incubated with both 500 nM of fluorescent lipid - Vybrant Dil (membrane probe) (Thermo-Fisher Scientific) and 500 nM of membrane-permeant (content probe) Green CMFDA cell tracker (Thermo-Fisher Scientific) in DMEM supplemented with Hoechst dye and incubated for 30 min.

Fusion pore detection also involved preparation of two separate population of cells at 75% confluency. In this case one population was incubated with 500 nM of membrane-permeant (content probe) Orange CMRA cell tracker (Thermo-Fisher Scientific) in DMEM with Hoechst dye, and the other population was incubated with 500 nM of membrane-permeant (content probe) Green CMFDA cell tracker (Thermo-Fisher Scientific) in DMEM with Hoechst dye and incubated for 30 min.

Subsequently, for each assay, the two separate populations were trypsinized and mixed in 1:1 ratio. The cell-mixture was replated on glass bottom dish in GM. Once cells attained 70% confluency, DM was added and used for quantifying non-fused, hemi-fused and fused cells at 2, 16, 24, 36, 48 hrs post DM addition.

Imaging was done via epifluorescence microscopy under 10X magnification. The number of nuclei that were in the particular stage of myogenesis was divided by the number of nuclei that are present in the field of image to calculate the percentage of cells of any particular phase phase. Fields with less than 70 cells were not used for these assays.

### Transfection

Cells were transfected with 1 μg of Gm7325 (Myomerger) (tGFP-tagged Mymx- Mouse predicted gene 7325 (Gm7325) transcript variant 3, OriGene - MG224234) plasmid DNA by using Lipofectamine 3000 (Invitrogen). Other treatments, if required, were performed 16 h after transfection. For the 2 hr condition, at 60% confluency, transfection was carried out in GM. 14 hrs post-transfection, DM was added to the cells such that 16 hrs post transfections (2hrs post DM) cells could be imaged. However, for rest of the time points/phases, cells were transfected in DM adjusting the time of transfection such that all imaging is done 14- 16 hrs post-transfection under every condition to minimize the effect of overloading cells with Myomerger. After 6 hours of the lipofectamine-based transfection media is replenished according to experimental requirements. As it is known that plasmid DNA mediated transfection reaches maximum level of expression at 24 hrs post transfection and too high concentration of Myomerger is detrimental for fusion, all experiments were performed within 14- 16hrs post transfection.

### Treatments

For quantifying the role of conical exogenous small lipids like Oleic acid (OA)(Oleic acid: Sigma-Aldrich) was dissolved in 30% ethanol and for quantifications of biophysical properties and fusion index, a working concentration of 10 µM was used. DM supplemented with OA was administered to cell as per experimental requirements with replenishment at every 24 hrs.

### Flipper-TR Fluorescence Lifetime imaging and quantification

For fluorescence lifetime imaging of Flipper-TR (Flipper TR membrane tension sensor-Cytoskeleton)(Colom et al., 2018), cells were treated with 1 µM of Flipper-TR for 45 min in DMEM (serum starved) and then washed with DM and imaged. The Abberior facility line built around the Olympus IX83 equipped with time correlated single photon counting (TCSPC module, PicoQuant) was employed to capture the Fluorescence lifetime of the probe. A 60x oil objective (1.42 NA) was used for the imaging keeping x, y pixel size as 200 nm. Imaging was done of the cell at nearly ∼ 1.5 – 2 µm above the basal membrane. Dwell time was set at 60 µs, excitation – 20%, bin time – 100 ps, bin count - 250.

For analysis, using MATLAB, the mean Fluorescence lifetime contributed by the peripheral plasma membrane of the cell is calculated per cell under different conditions. Membrane regions having flipper-TR probe intensity values more than 250 a.u. was considered for analysis of lifetime.

### siRNA knockdown treatment

For Myomerger knock-down assay, C2C12 cells were transfected with siRNA against Myomerger mRNA (IDT, mm.Ri.Gm7325.13.2) using lipofectamine-3000 (Invitrogen). When C2C12 cells were 50 % confluent in tissue culture(T-25) flask the first dose of 10 nM of siRNA was administered using protocol of transfection. After 48 hrs a second booster of 10 nM of siRNA was added to replenished GM and allowed to frow for the next 24 hrs (72 hrs after first dose). After 72h of incubation, cells were passaged to be re-seeded in imaging customised glass bottom dishes. After attaining 60 % confluency (which takes nearly 24 hrs) DM was added along with another booster of siRNA with timely changing (every 48h) of media supplemented with siRNA. The customized sequence of oligo duplex siRNA from IDT was given below:

Sense: 5’ AUCAAGGUCAGUUAAGAGAUGAUGT 3’
Anti-sense: 5’ ACAUCAUCUCUUAACUGACCUUGAUAC 3’

### Interference Reflection Microscopy

Interference Reflection Microscopy (IRM) was performed using Eclipse Ti-E motorized inverted microscope (Nikon, Tokyo, Japan) provided with adjustable field and aperture diaphragms, 60x Plan Apo (NA 1.22, water immersion) with 1.5x external magnification. sCMOS (ORCA Flash 4.0 Hamamatsu, Japan) camera was used for image acquisition in IRM, epifluorescence and DIC modes. For IRM, the system was equipped with a 100 W mercury arc lamp, an (546 ± 12 nm) interference filter with a 50:50 beam splitter. Fast time-lapse images of cells were acquired at 50 ms at a frame rate of 20 frames per second for 2048 frames(Biswas et al., 2017; Limozin & Sengupta, 2009).

#### Analysis of membrane height fluctuations

Calibration for determining the correlation of relative height of the membrane from the substrate to its corresponding intensity is done using 60 µm diameter polystyrene NIST beads (Bangs Laboratories Inc.). Cells were imaged in 3 ml of DM. Proper adherence of the cells to the glass substrate was ensured for selecting them for analysis. Further, from these selected adhered cells only regions that were within a range of ∼100 nm from the coverslip – also called the first branch region (FBR) were chosen and the size of these chosen FBRs were 12 × 12 pixels corresponding to ∼ 72 nm x 72 nm per pixel for IRM images and for the IRM images of the TIRF-IRM sets corresponded to corresponding to ∼ 65 nm x 65 nm per pixel. To maintain stringency in selecting the subset of regions for extracting parameters and estimating the membrane height fluctuations, the methodology designed previously (Biswas et al., 2017, 2019) has been followed.

For analysing the mechanical parameters of the cells under these two conditions over the time points, the power spectral density (PSD) was estimated by autoregression technique (probability of covariance or pcov) (MATLAB, Mathworks Inc, USA). For estimating active temperature (A), effective cytoplasmic viscosity (η_eff_), confinement (γ) and membrane tension (σ), PSD is fitted to the Helfrich-based Theoretical Model(Biswas et al., 2017):

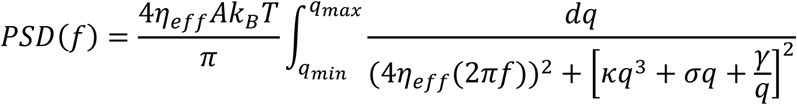

The fitting was done using MATLAB and fits with *R*^2^ > 0.9 were considered (Biswas et al., 2017) and the bending rigidity is kept constant at 15 k_B_T. The standard deviation of the membrane height over time series of a single pixel is estimated and averaged over single FBRs to obtain the membrane height fluctuations termed as SD_time_. Generally, effective membrane tension is termed as the fluctuation tension as the interpretation is dependent on the framework of membrane fluctuations(Shiba et al., 2016).

The pixel-wise standard deviation of the membrane height (SD_time_) over time series was also mapped.

### Sequential Correlative Total Internal Reflection Fluorescence (TIRF) - IRM microscopy: Local membrane mechanical properties

For correlative TIRF-IRM sequential imaging, Eclipse Ti-E motorized inverted microscope (Nikon, Tokyo, Japan) equipped with 100x - 1.4 NA water-immersion objective and sCMOS (ORCA Flash 4.0 Hamamatsu, Japan) was used. For IRM-based imaging a 100W Hg arc lamp with necessary interference filters (546 ± 12 nm), 50-50 beam splitter was used. Coherent OBIS laser of 488 nm was used for TIRF imaging.

The sequence followed for correlative TIRF-IRM sequential imaging for any field that had a fluorescent signal was TIRF-IRM-TIRF. For IRM imaging, time-lapse images at an interval of 50 ms with 20 frames per second rate were captured. For a particular field 2048 frames were captured for effective membrane fluctuation tension measurement. For TIRF-Imaging, images were acquired at an interval of 300 ms for 300 frames of the same field as that of IRM with a penetration depth of approximately 90-100 nm. TIRF movies captured before and after the IRM movie were used to verify the stability of the observed fluorescent pattern over the 1-2 min imaging period.

### Analysis of local membrane mechanistic properties by correlative TIRF-IRM

The first frame of the TIRF movie obtained after the IRM movie was used to obtain the pixel-wise correlation of local intensity with pixel-wise SD_time_ at the sites where fluorescently labelled Myomerger clusters at the plasma membrane. The IRM movies for every time points were template matched (using Fiji ImageJ software) with respect to time points as well as the fluorescence fields. Using the TIRF movies the fixed clusters were characterised.

All analyses were performed in MATLAB (MathWorks, USA) using experimentally acquired fluorescence images and corresponding pixel-wise mean membrane relative-height maps and SD_time_ maps generated for 2048 frames.

#### Curvature Extraction and Noise Sensitivity Analysis

All analyses were performed in MATLAB (MathWorks, USA) to quantify local membrane curvature of basal membrane at Myomerger clusters (from Fluorescence images) from experimentally derived mean relative height maps over 2048 frames and to evaluate robustness under simulated noise.

#### Image-based curvature analysis

Mean relative height maps and corresponding fluorescence images were loaded for each dataset. Objects were segmented from the fluorescence image using Gaussian smoothing (σ = 10 pixels) followed by background subtraction and intensity thresholding. The binary mask was refined using morphological operations (hole filling, erosion, dilation, and removal of small objects < 6 pixels), and connected components were labelled to identify individual structures. Morphological and intensity features were extracted using region properties, including centroid, weighted centroid, mean intensity, eccentricity, and equivalent diameter. Fixed objects were then selected manually from the labelled image, and the nearest segmented centroid was used as the reference point for each selected structure.

For each object, a local membrane patch of size 5×5 pixels were extracted from the relative height map, centred at the weighted centroid of Myomerger clusters. Boundary-touching objects were excluded. Each patch was mean-centred to remove offset variations.

#### Quadratic surface fitting and curvature estimation

Each local membrane patch was modelled using a second-order polynomial surface for getting membrane curvature at the weighted centroid of the Myomerger cluster (Bouzos et al., 2026):

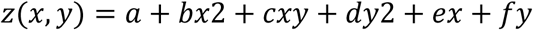

where coefficients were estimated using least-squares regression on the basis functions.[1, *x*2, *xy*, *y*2, *x*, *y*].Spatial coordinates were converted to nanometers using a pixel size of 65 nm.

The curvature was obtained from the Hessian of the fitted surface, with b,c,d as the coefficients obtained from quadratic fit.

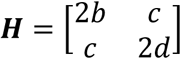

Principal curvatures (k_1_,k_2_) were computed as the eigenvalues by eigenvalue decomposition of the Hessian. k_1_, k_2_ correspond to the maximum and minimum local curvatures along two orthogonal principal directions on the membrane surface. Mean curvature (H) and Gaussian curvature (K) were defined as:

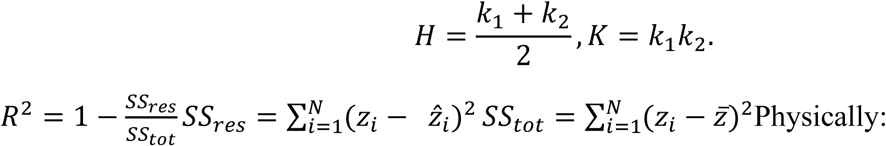

A) Both eigenvalues positive (K - +ve) indicate locally concave (exvagination - like) curvature.
B) Both eigenvalues negative (K - +ve) indicate locally convex (invagination - like) curvature.
C) Eigenvalues with opposite signs (K - -ve) indicate saddle-shaped geometry.

For visualization, a larger (31x 31) membrane patch centered on the selected object was extracted from the original relative height map. The larger patch was mean centred using the mean value of the fitted (5×5) local patch to maintain consistency with the curvature fitting procedure. The fitted (5×5) quadratic surface was then embedded at the center of the (31×31) region. The experimental membrane topography was rendered as a semi-transparent surface, while the fitted quadratic surface was overlaid as a mesh representation, enabling qualitative comparison between the measured membrane shape and the local curvature fit.

#### Local radial profile of relative height and membrane fluctautions

For each class of curvature state, local 31×31 patches centred on those corresponding object-weighted centroid were extracted from the fluorescence intensity image, membrane relative-height map, SD_time_ map.

To characterize radial membrane symmetry, line scans (length – 31 pixels, width – 1 pixel) passing through the patch center were generated along four fixed angular directions separated by 45°. The scan orientations corresponded to: 0°, 45°, 90°, 135°. For each direction, intensity, membrane height, SD_time_ values were sampled continuously along the line passing through the patch center across the full patch diameter. Spatial interpolation was used to estimate values at non-integer pixel coordinates along each radial direction. The directional line profiles obtained from the four orientations were subsequently averaged to generate a rotationally averaged radial profile for each object:

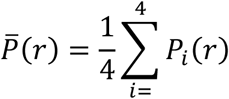

where P_i_(r) represents the profile measured along the i^th^ angular direction. To reduce directional asymmetry and local noise, radial profiles were further symmetrized about the patch center by averaging values located at equal distances on opposite sides of the center position. This produced symmetric radial representations of membrane intensity, relative height, and SD_time_ distributions surrounding each object.

For each distance d from the center, values located symmetrically about the center position were averaged: 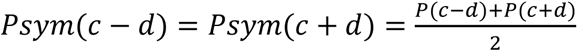, where c denotes the center index of the radial profile.

For each object, the mean membrane height within the 31×31 patch was calculated and subtracted from the radial height profile and SD_time_ profiles, for quantification. Objects were classified according to the mean substracted height value of line scans at the object centroid.

For each object of each curvature class, radial scan directions were visualised on the local membrane patches. Two-dimensional membrane height maps and three-dimensional surface renderings were generated for qualitative inspection.

From all the objects under each category the symmetric line scan profiles for relative height, SD_time_ centered around the cluster centroid were averaged and the median profile with corresponding s.e.m for each category was plotted. Only for better representation purposes both negative and positive distance were plotted – however the data is same for two points at same distance from the centre. The difference in relative height and SD_time_ of the membrane at the cluster centre from that at the edges of the clusters (∼1 µm) is denoted as the ΔH and ΔSD_time_. This value gives a quantification of the depth of the local membrane bending and the change in local membrane fluctuation pattern due to Myomerger clustering. Mann–Whitney U test was performed to calculate the statistical significance of the various mechanical properties at various distance from the centre. All plots were done using OriginPro software.

#### Transition State analysis

Raw IRM intensities for 2048 frames were converted to membrane height using bead calibration parameters. To improve the signal-to-noise ratio, every 50 consecutive frames were averaged to generate a single height map for subsequent analysis.

Objects were segmented from TIRF images using Gaussian background subtraction, intensity thresholding, hole filling, and morphological filtering. Individual objects were manually selected, and the weighted centroid of each object was used to extract a local region of interest.

For each averaged frame, a 5 × 5 pixel patch centered on the object was fitted with a second-order quadratic surface using least-squares regression. The fitted surface was used to compute the Hessian matrix, from which the principal curvatures (k₁ and k₂), mean curvature (H), and Gaussian curvature (K) were calculated. Based on the signs of the principal curvatures, membrane topology was classified as Peak (k₁ < 0 and k₂ < 0), Valley (k₁ > 0 and k₂ > 0), or Saddle (principal curvatures of opposite sign). The quality of the quadratic fit was assessed using the coefficient of determination (R^2^).

Morphological transitions were quantified by tracking the sequence of curvature states in consecutive time points for each object across time. For every pair of consecutive frames, transitions between the three states (Peak, Saddle, and Valley) were counted to generate a 3 × 3 transition matrix, where each matrix element represented the frequency of transitions from one state to another. Transition counts were normalized to obtain transition probabilities. To assess whether the observed transition dynamics were non-random, state sequences were randomly shuffled while preserving the overall state composition, and transition probability matrices were recalculated. Observed and shuffled transition probabilities were compared at both the object and file levels using the Mann-Whitney U test. Additionally, these values were compared with random non Myg cluster regions from the same cells (regions that donot have any fluorescence intensity.

### Stimulated Emission Depletion microscopy technique (STED)

#### Sample preparation (fixed)

After successful fixation of the cells using protocol earlier described, the coverslips containing the cells were subjected to the 3 hrs of blocking using 0.2% gelation. Then cells were incubated with primary antibody against mouse Myomerger exoplasmic domain - ESGP polyclonal antibody (Invitrogen, PA5-47639 - raised in sheep) at 1:200 dilution in 0.2% Gelatin for 2 hr at room temperature. For Myomaker and Clathrin co-staining, the primary antibody against Myomaker (Anti-TMEM8C antibody (ab188300)) and Clathrin (Clathrin heavy chain Rabbit mAb (Abclonal – human antigen) at 1:200 respectively, in 0.2% gelatin, were added for 2 hr at room temperature together with Myomerger primary antibody. After incubation, the primary antibodies were washed thoroughly with 1X PBS. The Donkey Anti-Sheep IgG abberior STAR RED secondary antibody (abberior) was added to cells at a 1:400 dilution in 0.2% gelatin along with Secondary antibodies for Myomaker (Goat Anti-Rabbit IgG H&L (Alexa Fluor® 568) (ab175471)) or Clathrin (Goat Anti-Rabbit IgG H&L (Alexa Fluor® 568) (ab175471)) at dilution of 1:200 and incubated for 1.5 hr. Following the staining, the sample was washed three times in 1X PBS. The samples that were stained for Actin were incubated with Alexa Flour 568 Phalloidin after its required antibody staining and incubated for 45 min, followed by a thorough was with 1X PBS and then mounted on a pre-cleaned grease-free glass slide with MOWIOL 4- 88 (Sigma Aldrich) mounting media (Ghosh et al., 2025).

2D – STED Imaging for visualising Actin cytoskeleton, Myomerger clustering and Myomaker – Myomerger colocalization studies

2D - STED Microscopy was performed using an inverted microscope (Abberior’s Facility Line STED) built around a 60x oil objective (1.42 NA) and Olympus IX83. To ensure proper alignment of the STED and confocal point spread function (PSF), an auto-alignment sample containing uniform-sized multifluorescent beads (∼120 nm) was used. A pulsed 775 nm laser was utilized for the depletion of red and far-red fluorescence. The point spread function (PSF) for 2D-STED has a diameter of ∼30 nm along the xy and ∼500 nm along z axes. Dwell time was set at 5 µs with 30% STED power and 15% excitation for imaging.

For 2D-STED imaging for fixed for actin fibres and Myomerger clusters in cells, a 1.0 AU pinhole was used with 20 nm as xy resolution and step size of 300 nm was standardised for actin cytoskeleton and Myomerger clustering visualization under different conditions. For quantification, the base of the cell (first focus) was considered to quantify the PM surface Myomerger expression and Actin distribution.

In the 2D-STED imaging process, a xy pixel size of 15 nm was used to collect the images of Myomaker and Myomerger clusters.

3D - STED Imaging for visualising Actin cytoskeleton, Myomerger cluster colocalization and Myomerger and Clathrin cluster co-distribution at the basal membrane.

3D - STED Microscopy was performed using an inverted microscope (Abberior’s Facility Line STED) built around a 60x oil objective (1.42 NA) and Olympus IX83. To ensure proper alignment of the STED and confocal point spread function (PSF), an auto-alignment sample containing uniform-sized multifluorescent beads (∼120 nm) was used. A pulsed 775 nm laser was used to deplete red and far-red fluorescence. The point spread function (PSF) for 3D-STED has a diameter of ∼70 nm along both the xy and z axes. This allows to restrict the signal accumulation from the basal membrane of cells, and thus the co-distribution of Myomerger clusters, actin cytoskeletal. Dwell time was set at 5 µs with 30% STED power and 15% excitation for imaging.

In the 3D-STED imaging process, a xy pixel size of 50 nm and a step-size along z of 70 nm was used to collect the images of actin fibres and Myomerger clusters.

### FLIM-STED live imaging

#### Sample preparation (live) for 2D - STED and Lifetime imaging

Live cells that are 70% confluent are administered with DM and kept for 24 hrs. After that, media is replaced using 1 ml of DMEM (37°C). Prior to replacing this 1ml of M1 media was mixed with 1:100 dilution of Myomerger exoplasmic domain - ESGP polyclonal antibody (Invitrogen, PA5-47639 - raised in sheep) primary antibody. Incubated for 30 min. after that it is thoroughly washed with M1 media and again 1:100 dilution of Donkey Anti-Sheep IgG Abberior STAR RED secondary antibody (Abberior) was prepared using DMEM and added to cells for 20- 30 min depending on cell health. Immediately after incubation cells are washed with DMEM and for fluorescence lifetime imaging of Flipper-TR (Flipper TR membrane tension sensor-Cytoskeleton)(Colom et al., 2018), cells were treated with 1 µM of Flipper-TR for 45 min in DMEM and then washed with DMEM media and replenished with DM for imaging.

#### Imaging

For 2D-STED imaging for live cells with Flipper-TR lifetime, a 1.0 AU pinhole was used with 50 nm as xy resolution was standardised. Because in this case FLIM and Sted were simultaneously done on live cells to optimize time duration of imaging and resolution xy pixel size was kept at 50 nm instead of 20 nm unlike fixed sample 2D STED. Images at two planes were captured – one at the base – first focus of Myomerger signal and other at 4-5 µm above the basal membrane to reach the midplane of the cell. Dwell time was set at 15 µs, excitation – 15%, STED power – 25%.

For FLIM imaging along with live cell STED, excitation – 20%, bin time – 100 ps, bin count – 250 was maintained.

#### Analysis

From the STED images of object detection was done to detect Myomerger clusters. From the distribution of Myomerger cluster, equivalent diameter the clusters that were < 400 nm diameter were considered as small clusters and > 400 nm are considered as laeger clusters. For detecting the change in lifetime around Myomerger cluster on the basal plane, line scans of 17 pixels length and 1 pixels width was performed in both x and y direction centred around cluster object centroids. Then the mean line scan was obtained. The profiles were also averaged based on distance from centre. Only for better representation purposes both negative and positive distance were plotted – however the data is same for two points at same distance from the centre. The FWHM from the STED images were calculated for gaussian fits of the intensity line scans from the cluster objects to get the size of the clusters. Analysis done using MATLAB and OriginPro software. For the change in lifetime around Myomerger cluster on the lateral membrane of the midplanes, manual line scans were performed of 15 pixels length and 1 pixel’s width and then averaged based on centre of cluster (with 0 representing cluster centre for all cases). FWHM of the gaussian fits of these line scans was used to derive the size of the clusters. This was primarily done in ImageJ and OriginPro software. Furthermore, for all classes small clusters, larger clusters and the clusters at the lateral membrane, 11 x 11 pixels patches with centroid as the patch centre is extracted. In this patch, the Spearman correlation coefficient of pixel-wise Myomerger cluster intensity with Flipper TR Fluorescence Lifetime. This analysis were done using MATLAB (MathWorks, USA). Mann–Whitney U test was performed to calculate the statistical significance at various distance from centre.

### Number density, fractional area coverage and morphological quantification of Myomerger clusters

To analyze the degree of Myomerger clustering STED/TIRF images were taken and these images were subjected to object detection in MATLAB. To create the binary images for analysis, a Gaussian blur operation was first performed, followed by subtracting it from the original image to reduce noise. The noise-reduced image was then converted to values between 0 and 1, and proper thresholding was performed to form a binary image which subsequently used for object detection. Alternatively, adaptive thresholding depending on signal to noise ratio of cluster was performed, for better foreground to background detection. Objects encompassing less than 4 pixels for STED (mean intensity threshold – 2.5) and 5 pixels for TIRF (size threshold) were rejected.

For quantification, the number of objects per unit area of the cell was calculated and termed as number density. The ratio of the area covered by the objects to the total area of the cell depicts the fractional area coverage. Further, the area (A) (number of pixels per object*(pixel size)^2^) and the mean Intensity of each object was also quantified. For each segmented object, the pixel area was converted to an equivalent circular diameter (d_eq_), 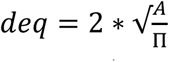, defined as the equivalent diameter of a circle with identical area. Measurements were subsequently converted from pixels to physical units (nm or µm) using the calibrated pixel size of the imaging system. The sizes referred in text depicts median as the value.

### Bodipy cholesterol co-staining with Myomerger

Cells at 70 % confluency were washed with M1 media. It was incubated with 1µM of Bodipy Cholesterol (stock 1 mM in PBS) in M1 media for 30 min at 37C and 5% CO2. Then the cells were fixed for immunostaining against Myomerger for STED imaging.

The Abberior facility line built around the Olympus IX83 equipped with 60x oil objective (1.42 NA) was used for the imaging keeping x, y pixel size as 15 nm. Imaging was done of the basal membrane (first focus) of the cell. Dwell time was set at 5 µs. Myomerger clusters were imaged by 2D STED and Cholesterol using simultaneous confocal scanning.

### Minimum distance from boundary of nearest neighbour analysis

For estimating the distance between F-actin/cholesterol nanodomains with Myomerger clusters object detection on both channels is performed. The perimeter of the cholesterol / F-actin objecta were extracted to define the cholesterol boundary, and the shortest Euclidean distance from the weighted centroid of each real and randomized Myg cluster to the nearest cholesterol/F-actin boundary pixel was calculated. Myomerger cluster randomization was done by Monte Carlo randomisation of the centroids coordinates of real objects within the cell boundary. Analysis was done in MATLAB and Plotting in OriginPro Software.

### Statistical tests

A Mann–Whitney U test was performed for statistical significance testing (ns denotes p > 0.05, * denotes p < 0.05, ** denotes p < 0.001). performed using OriginPro software. For all box plots, the error is SD and for all line plots, the error is SEM unless mentioned otherwise. All normalised distribution plots and violin plots have the corresponding mean of the distribution marked and were generated using MATLAB.

## RESOURCE AVAILABILITY

### Lead Contact

Further information and request for resources and reagents should be directed to and will be fulfilled by the lead contact, Bidisha Sinha.

### Materials Availability

All new materials and methods generated in this study will be available upon request from Bidisha Sinha.

### Data and code availability

Data and custom codes used in this study for analysis will be available upon request to the lead contact.

### Author contribution

BS conceptualized the project. MC and UM set up the model system. All experiments were performed by UM. UM analysed all the experiments. BS and UM wrote the first draft of the manuscript and prepared the display items. All authors edited the manuscript. BS acquired the funding for the work.

## Acknowledgements

B.S. acknowledges support from Wellcome Trust/DBT India Alliance fellowship (grant number IA/I/13/1/500885), SERB (grant number SERB_CRG_2336) U.M. thanks IISERK for providing her fellowship. MC thanks CSIR for their fellowship. The authors are also thankful to the Builder Imaging facility (BT/INF/22/SP45383/2022) for super-resolution STED imaging.

