## Supplementary Material for "Size-dependent membrane remodelling at Myomerger clusters during myoblast fusion"

**Supplementary Text Material:**

Extended methods:

1. Area of objects = the total number of pixels in the object multiplied by the area of each pixel
2. Equivalent diameter = $2\sqrt{\frac{A}{\pi}}$ where A is the area per object .
3. Number Density = $\frac{N}{Ar}$ where N is the number of objects detected from the image. "Ar" denotes the Area of the ROI for a particular z in which the objects are detected.
4. Fractional Area Coverage = $\frac{\Sigma iAi}{Ac}$ where $A$ represents the area of each object while Ac represents the area of the ROI

#### M1 media preparation

M1 media composed of 150 mM NaCl (Sigma-Aldrich), 1 mM $MgCl_2$ (Merck), 20 mM HEPES (Sigma)

#### Fusion Index calculation

For measuring fusion index, after differentiation (appearance of myotubes – 72 hrs in DM), cells were fixed following the previous protocol. MyHC (Myosin Heavy chain) is a known differentiation marker (Chakraborty et al., 2022). After fixation, the cells are treated with 0.2% Triton-100X for permeabilisation for 2 min at RT. Subsequently, cells were incubated with 3 ml of 0.2% gelatin (Sigma-Aldrich) solution as blocking agent, for 3 h at room temperature (RT). Primary antibody treatment was done with Myosin 4 Monoclonal Antibody – Anti-MyHC antibody (MF20, Invitrogen, eBioscience) staining (raised against mouse) is carried out at 1:200 dilution and incubated at 4°C overnight. After washing the cells with PBS, goat anti-mouse IgG H&L, Alexa Fluor 488 secondary antibody (Abcam, ab150117) at a dilution of 1:500 was added and kept for 2 h at RT. After incubation, it was washed with PBS and then stained with DAPI (1:1000).

MF20 and DAPI- labelled myotubes were imaged via epifluorescence microscopy and quantified for fusion index calculations. The fusion index was estimated by dividing the number of nuclei in a frame as a part of Myotube(s) by the total number of nuclei in the same frame. The fraction obtained was multiplied by 100 for calculating the percentage.

**Goodness of Fit for curvature analysis**

Goodness of fit was quantified using the coefficient of determination:

$$R^2 = 1 - \frac{SS_{res}}{SS_{tot}}$$

where the residual sum of squares is defined as

$$SS_{res} = \sum_{i=1}^{N} (z_i - \hat{z}_i)^2$$

and the total sum of squares is

$$SS_{tot} = \sum_{i=1}^{N} (z_i - \bar{z})^2$$

Here, $z_i$ represents the observed membrane height at point i, $\hat{z}_i$ is the fitted surface height obtained from the quadratic model, $\bar{z}$ is the mean of the observed heights, and N is the total number of points in the local membrane patch. Each object was classified based on curvature signs: negative–negative eigenvalues corresponded to peaks, positive–positive to valleys, and mixed signs to saddle structures. All object-level metrics (curvatures, $R^2$, intensity, size, centroid) were stored for statistical analysis after classifying them into peak (internally indented) , valley (externally bulged) and saddles based on the eigenvalues. A threshold of 0.7 as goodness of fit was considered for each curvature state.

**Noise sensitivity analysis**

To assess robustness of curvature estimation, synthetic membrane surfaces were generated using the same quadratic model with known curvature parameters. Two cases were considered: peak and valley geometries with equal principal curvatures ($k_1 = k_2 = \pm 10^{-5}$ $nm^{-1}$). Corresponding theoretical mean and Gaussian curvatures were computed analytically. Additive Gaussian noise with standard deviations $\sigma = [0,0.1,0.2,0.5,1,2,5,10]$ nm was introduced to the surface. For each noise level, 20 independent realisations were generated. Each noisy surface was refitted using the same quadratic regression framework, and curvatures were re-estimated from the Hessian eigenvalues. Performance was evaluated using: coefficient of determination ($R^2$), recovered mean curvature (H), and recovered Gaussian curvature (K). For each noise level, mean and standard deviation were computed across repetitions to quantify stability and noise sensitivity of the curvature extraction method.

**Convolution lookup table**

A MATLAB simulation was performed to evaluate how Gaussian blurring affects the measured diameter of circular objects. Binary circular objects with radii ranging from 2 to 200 pixels (0.5-pixel increments) were generated and convolved with a normalized two-dimensional Gaussian point spread function (PSF) having a full width at half maximum (FWHM) of 30 pixels. The resulting images were normalized and segmented using Otsu's thresholding method, followed by hole filling and removal of small, isolated regions. The largest connected object was retained, and its equivalent circular diameter was measured using MATLAB's regionprops function. This procedure was repeated for all object sizes to obtain the relationship between the original object diameter and the measured diameter after Gaussian blurring

**Total internal reflection fluorescence microscopy (TIRF) and epifluorescence microscopy**

For TIRF, Olympus IX-83 inverted microscope (Olympus, Meliville, NY) equipped with a 100 × 1.49 NA oil immersion TIRF objective (PlanApo, Olympus) was implemented. A CMOS camera (ORCA Flash 4.0 Hamamatsu, Japan) was used for image acquisition. A 488-nm laser beam was used as a laser source for TIRF. All images were captured at 300 ms exposure time and ~100 nm penetration depth. For time series, for classification of stuck clusters, images were acquired for 600 frames at 300 ms (3 min time series). For TIRF pixel size is 65nm. All the surface expression analysis was done using TIRF images, unless mentioned otherwise.

For epifluorescence, a mercury arc lamp was used, and images were taken using FITC, TRITC and DAPI filters with ×60 objectives. All snapshots were acquired at 300 ms of exposure time. For epifluorescence, pixel size was 108 nm. All percentage of cell counts were performed by epifluorescence-based images.

**Immunofluorescence for TIRF imaging**

For immunofluorescence, C2C12 cells, at different stages of differentiation, were first washed of its culture media (GM/DM) properly with phosphate-buffered saline (PBS, Sigma-Aldrich) and were fixed with 4% paraformaldehyde (Sigma-Aldrich) for 15 min followed by a thorough wash with PBS. Cells were then incubated in 0.1 M glycine (Sigma-Aldrich) for 5 min followed by another wash with PBS. Subsequently, cells were incubated with 3 ml of 0.2% gelatin (Sigma-Aldrich) solution as blocking agent, for 3 h at room temperature (RT).

For quantification of surface level Myomerger by TIRF, Primary antibody treatment (against Myomerger) was carried out with ESGP polyclonal antibody (Invitrogen, PA5-47639 - raised in sheep) at 1:200 dilution in 0.2% gelatin and kept at 4°C overnight. After washing the cells with PBS, Donkey anti-sheep Alexa Fluor 488 secondary antibody (Invitrogen, A-11015) at a dilution of 1:500 was added and kept for 2 h at RT. DAPI (Sigma-Aldrich) with 1:1000 dilution was used for nucleus visualization. Cells were washed after each step with PBS, and fixed cell imaging was carried out finally in 2 ml of PBS.

**List of Supplementary figures:**

S1 - Myomaker is involved in hemifusion and Myomerger is involved in fusion pore formation

S2 - Temporal alteration in membrane fluctuations and tension profile during differentiation

S3 - Effect of $siRNA_{myg}$ treatment on overall abundance of Myomerger in cells and its impact on fusion index

S4 - Myomaker increases effective basal membrane tension in both early and late phase of fusion

S5 - Myomerger clusters on basal membrane

S6 – Schematic optical Ray diagram of Sequential TIRF- IRM microscope system

S7 - Myomerger clusters significantly alter local membrane topology

S8- Smaller pool of Myomerger cluster dampens membrane fluctuation at the centre, and larger ones are mostly association invaginations

S9 – Video of a patch of membrane that has net average IN topology for 100s.

S10 - Video of a patch of membrane that has net average EX topology for 100s.

**Figure S1**

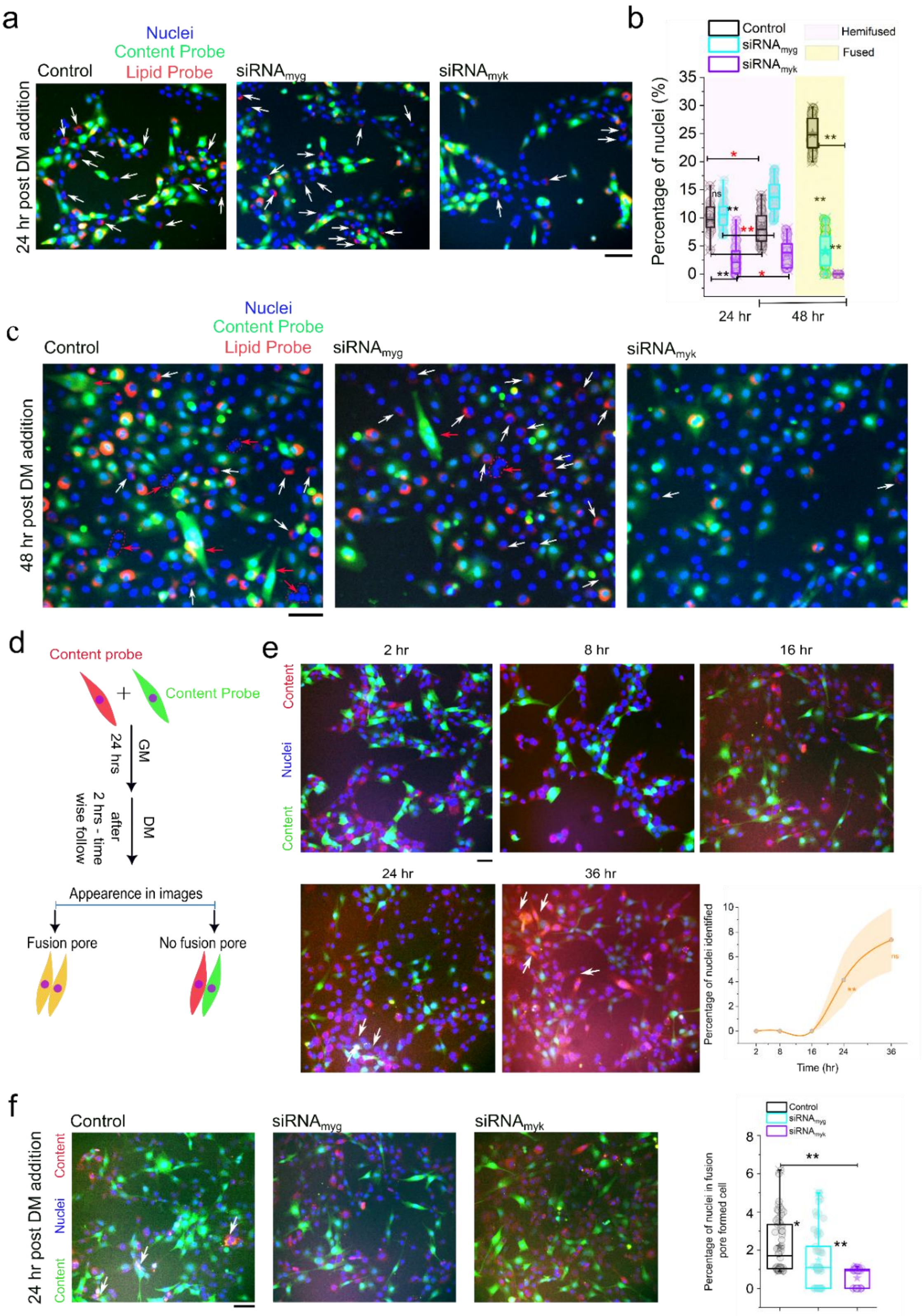

a
Nuclei
Content Probe
Lipid Probe
Control
siRNAmyg
siRNAmyk
24 hr post DM addition
b
Control
siRNAmyg
siRNAmyk
Hemifused
Fused
Percentage of nuclei (%)
ns
24 hr
48 hr
c
Nuclei
Content Probe
Lipid Probe
Control
siRNAmyg
siRNAmyk
48 hr post DM addition
d
Content probe
Content Probe
GM
24 hrs
DM
after
2 hrs - time
wise follow
Appearence in images
Fusion pore
No fusion pore
e
2 hr
8 hr
16 hr
24 hr
36 hr
Content
Nuclei
Content
Percentage of nuclei identified
Time (hr)
f
Control
siRNAmyg
siRNAmyk
24 hr post DM addition
Content
Nuclei
Content
Percentage of nuclei in fusion pore formed cell

**Figure S1 – Myomaker is involved in hemifusion and Myomerger is involved in fusion pore formation**

(a,c) Representative (green – content probe, red - lipid probe, blue – (DAPI-stained) nuclei) epifluorescence images to quantify the percentage of hemi-fused cells after 24 hrs and 48 hrs of DM addition for control Myomerger knockdown ($siRNA_{myg}$) and Myomaker knockdown ($siRNA_{myk}$)conditions. White arrows mark the hemi-fused cells, and red arrows mark the fused cells. (b) The Percentage of hemifused and fused cells at 24 hrs and 48 hrs post DM addition under Control (black), $siRNA_{myg}$ (cyan), $siRNA_{myk}$ (purple). The magenta shaded region marks the mono-nucleated cells of 24 hrs, and 48 hrs post DM addition states and yellow shaded region shows the fused population observation post 48 hrs of DM. $N_{field}$ - Control – 51, $siRNA_{myg}$ – 87, $siRNA_{myk}$– 38 (d) The schematic for differentially labelling the C2C12 cells for identification of fusion pore formation. (e) The representative (green – content probe, red – content probe, blue – (DAPI-stained) nuclei) epifluorescence images to quantify the percentage of fusion pore formed cell sthat show content mixing after 2 hrs,8hr, 16 hr, 24 hr, 36 hr of DM addition. White arrows mark the dual colour volume labelled cells – marking fusion pore formation. Represents the temporal change in percentage of nuclei in population of cells showing content missing post DM addition. The shaded region marks the SEM. Data are median ± s.e.m. $N_{field}$ - 2 hr – 56, 8 hr – 59, 16 hr – 69, 24 hr – 85, 36 hr – 42. (f) Representative (green – content probe, red - content probe, blue – (DAPI-stained) nuclei) epifluorescence images to quantify the percentage of fusion pore formed pairs of cells after 24 hrs of DM addition for control Myomerger knockdown ($siRNA_{myg}$) and Myomaker knockdown ($siRNA_{myk}$)conditions. White arrows mark the content mixed dual labelled cells. The Percentage of dual colour volume labelled cells at 24 hrs post DM addition under Control (black), $siRNA_{myg}$ (cyan), $siRNA_{myk}$ (purple). $N_{field}$ - Control – 67, $siRNA_{myg}$ – 58, $siRNA_{myk}$– 42. The Mann-Whitney U-test was used for analysis. * $p < 0.05$, ** $p < 0.001$, ns $p > 0.05$. For all line plots, the median line profile with s.e.m as the shaded region is depicted. For all box plots, error is SD.  Scale bar – 10 μm.

#### Figure S2

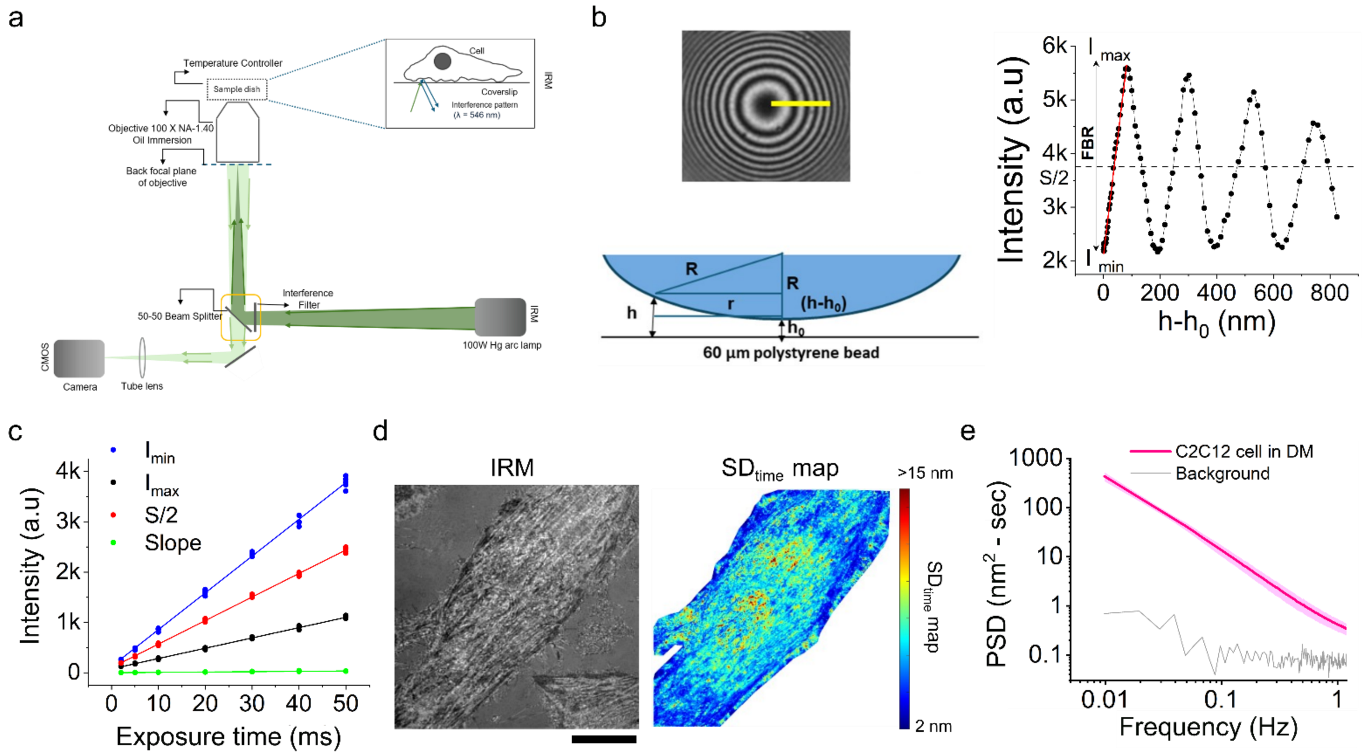


**Figure S2- Temporal alteration in membrane fluctuations and tension profile during differentiation –**

(a) Schematic to explain the optical illumination path used to achieve Interference Reflection microscopy (IRM). (b) (left panel) IRM images of a 60 µm polystyrene bead attached on the glass surface showing interference pattern for calibration Intensity with relative height. Cross-sectional representation of bead attached on the glass surface displaying the profile of its distance from coverslip (height with $h_0$ being the reference and h- $h_0$ used as relative height. (right panel) Intensity vs relative height profile derived from the bead in left panel along the yellow line. Intensity minima ($I_{min}$), maxima ($I_{max}$) and S/2 (background intensity) were marked out. (c) Typical image of basal membrane of cell imaged using IRM with corresponding pixel-wise $SD_{time}$ map. (d) Dependence of $I_{min}$ , $I_{max}$ and S/2 on exposure times (e) Fitted PSD (red bold median line – error shaded is s.e.m) for few FBR along with a background region indicated. (Biswas, Alex and Sinha, 2017).

**Figure S3**

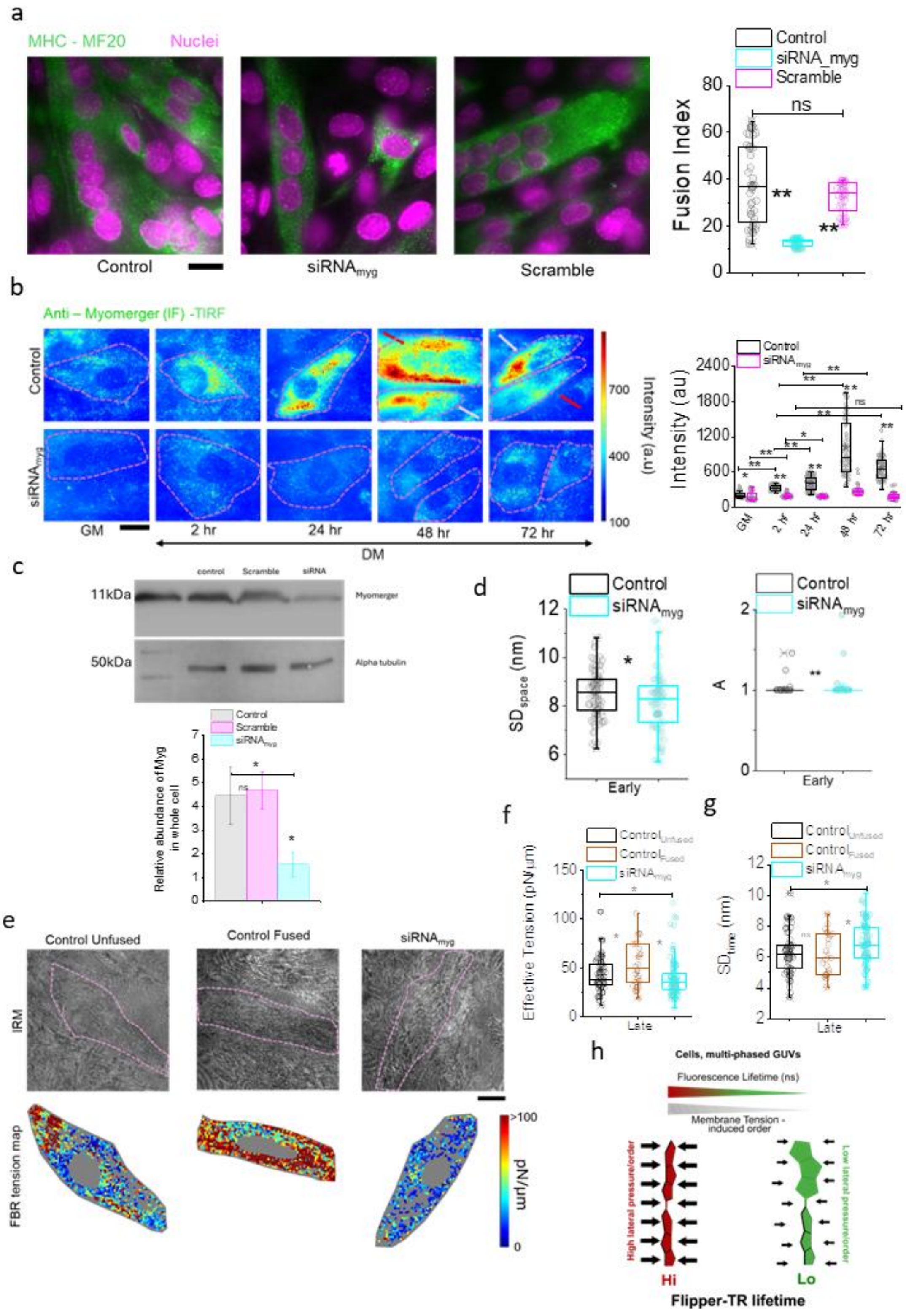
a
MHC - MF20
Nuclei
Control
siRNA_myg
Scramble
Fusion Index
ns
b
Anti – Myomerger (IF) -TIRF
Control
GM
2 hr
24 hr
48 hr
72 hr
DM
Intensity (a.u)
Intensity (au)
c
control
Scramble
siRNA
11kDa
50kDa
Myomerger
Alpha tubulin
Relative abundance of Myg in whole cell
d
Control
Early
e
Control Unfused
Control Fused
IRM
FBR tension map
pN/µm
f
Effective Tension (pN/µm)
Late
g
h
Cells, multi-phased GUVs
Fluorescence Lifetime (ns)
Membrane Tension - induced order
High lateral pressure/order
Low lateral pressure/order
Hi
Lo
Flipper-TR lifetime

**Figure S3– Effect of $siRNA_{myg}$ treatment on overall abundance of Myomerger in cells and its impact on fusion index**

(a)(left)Representative Immunofluorescence (IF) (epifluorescence) of MF20 and DAPI staining for control, $siRNA_{myg}$, and Scramble-treated cells after 72 hrs of exposure to DM. (right) Fusion index and number of nuclei from each myotube under each condition are compared. $N_{repeat}$ – 3, $N_{field}$ – control (49 – black), $siRNA_{myg}$ (32 - cyan), Scramble (37 - magenta). (b) (left) Representative TIRF images of time point wise surface enhancement of Myomerger content of C2C12 cells and its downregulation on Myomerger knockdown. Cell boundary marked by magenta dotted line. Red arrows mark fused cells and white arrows mark non-fused cells. (right) Temporal profile of cell-wise surface enhancement of Myomerger at the basal membrane and its comparison with Myomerger knockdown conditions. (c) Representative blot of Myomerger abundance at 24 hrs and its depletion by knockdown. The relative abundance of Myomerger on $siRNA_{myg}$ treatment in whole cell. (d) The comparative study of $SD_{space}$ and Activity of basal membrane of cells on Myomerger knockdown. (e) Representative IRM images with corresponding FBR tension maps for late phase cells under control, Myomerger knockdown for both mono nucleated and fused cells. (f) The comparative study of effective tension of basal membrane of cells on Myomerger knockdown. (g) The comparative study of $SD_{time}$ basal membrane of cells on Myomerger knockdown. $N_{repeat}$ – 3, $N_{ell}$ – control unfused (51– black), $siRNA_{myg}$ (55 - cyan), control fused (25 - brown). (g) The Schematic of the mode of action of tension and lipid compaction sensor probe Flipper-TR. The Mann-Whitney U-test was used for analysis. * $p < 0.05$, ** $p < 0.001$, ns $p > 0.05$. For all box plots, error is SD. Scale bar – 10 µm.

**Figure S4**

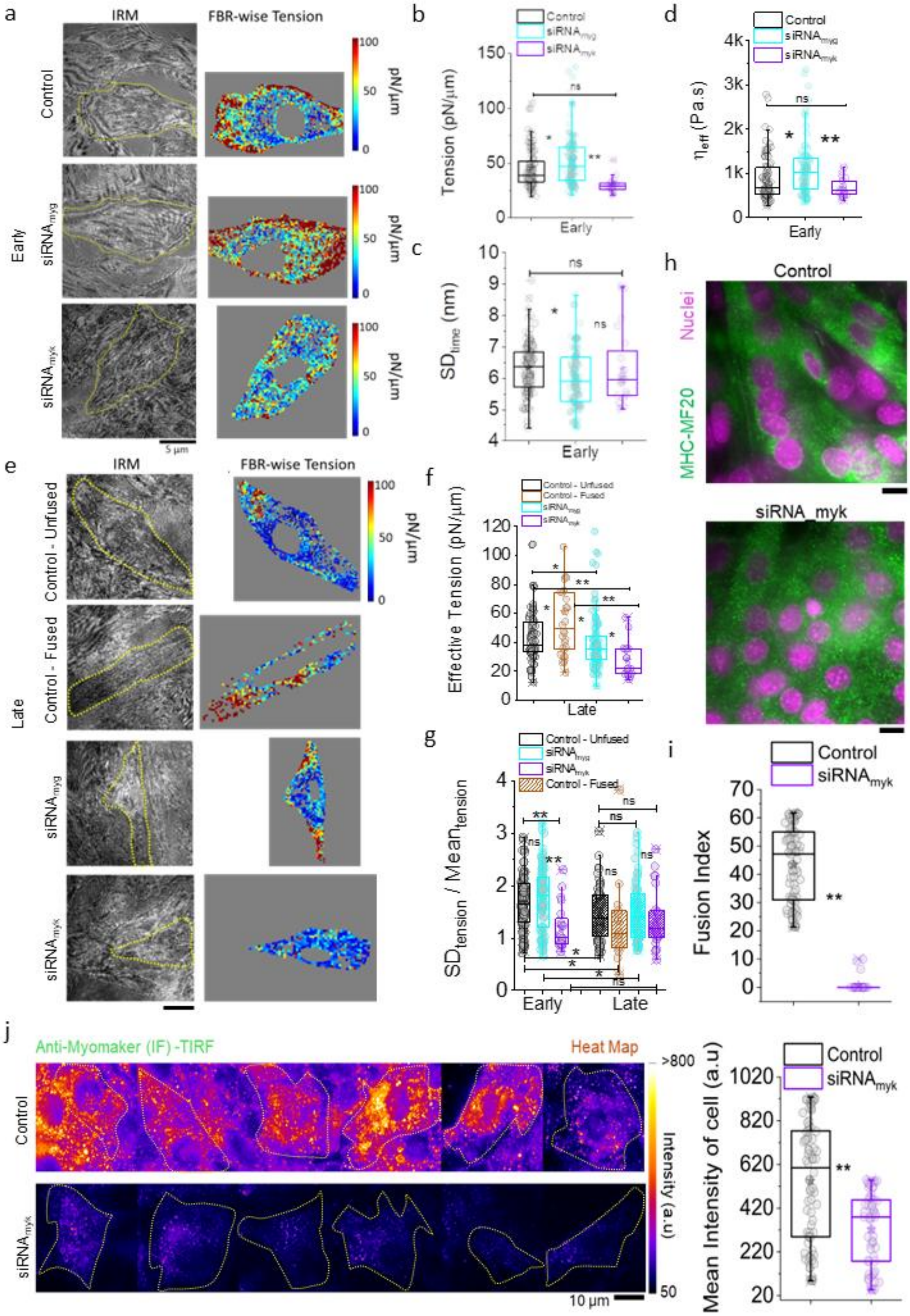
a
IRM
FBR-wise Tension
Control
Early
siRNA_myg
siRNA_myk
pN/µm
100
50
0
5 µm
b
Control
siRNA_myg
siRNA_myk
Tension (pN/µm)
Early
c
SD_time (nm)
Early
d
η_eff (Pa.s)
Early
e
IRM
FBR-wise Tension
Control - Unfused
Control - Fused
Late
f
Control - Unfused
Control - Fused
Effective Tension (pN/µm)
Late
g
SD_tension / Mean_tension
Early
Late
h
Control
Nuclei
MHC-MF20
siRNA_myk
i
Fusion Index
j
Anti-Myomaker (IF) -TIRF
Heat Map
>800
Intensity (a.u)
50
10 µm
Mean Intensity of cell (a.u)

**Figure S4– Myomaker increases effective basal membrane tension in both early and late phase of fusion**

(a) Representative IRM images with corresponding FBR tension maps for early phase (24 hrs) C2C12 cells that were either treated with $siRNA_{myg}/siRNA_{myk}$. (b) Comparative effective tension profiles of early phase cells on Myomerger and Myomaker knockdown condition. (c) Comparative $SD_{time}$ profiles of early phase cells on Myomerger and Myomaker knockdown condition. (d) Comparative effective viscosity of early phase cells on Myomerger and Myomaker knockdown condition (e) Representative IRM images with corresponding FBR tension maps for late phase (48 hrs) C2C12 cells that were either treated with $siRNA_{myg}/siRNA_{myk}$ (f) Comparative effective tension profiles of early phase cells on Myomerger and Myomaker knockdown condition. (g) Comparative $SD_{tension}/Mean_{tension}$ profiles of early and late phase cells on Myomerger and Myomaker knockdown condition. (h) Representative Immunofluorescence (IF) (epifluorescence) of MF20 and DAPI staining for control, $siRNA_{myk}$, and Scramble-treated cells after 72 hrs of exposure to DM. (i) Fusion index and number of nuclei from each myotube under each condition are compared. (j) Representative TIRF images 24 hr post DM treated cells surface eexpression of Myomaker content of C2C12 cells and its downregulation on Myomaker. Cell boundary marked by yellow dotted line. (right) Comparative profile of cell-wise surface expression of Myomaker at the basal membrane and its comparison with Myomaker knockdown conditions. $N_{repeat}$ – 3. The Mann-Whitney U-test was used for analysis. * $p < 0.05$, ** $p < 0.001$, ns $p > 0.05$. For all box plots, error is SD. Scale bar – 10 μm.

**Figure S5**

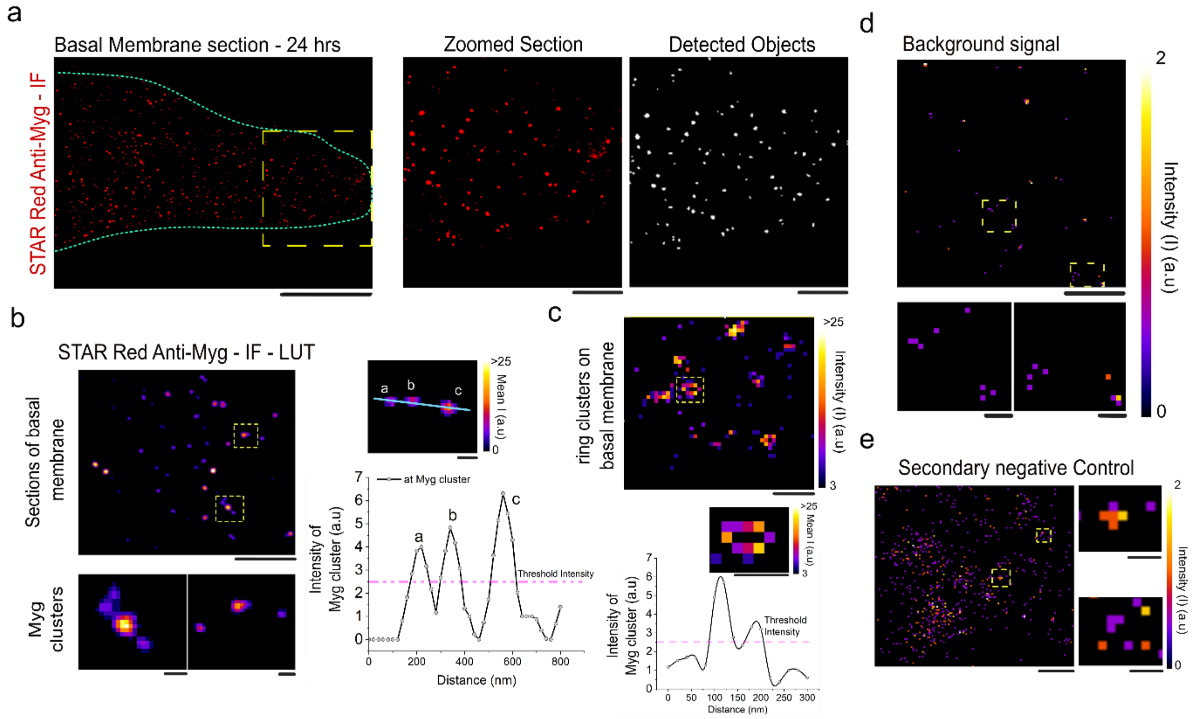


**Figure S5– Myomerger clusters on basal membrane**

(a)Representative 2D STED images of C2C12 cells taken 24 hours after DM treatment display immunofluorescence (Abberior STAR Red) against the endogenous Myomerger protein. A cyan dotted line indicates a section of the cell's basal membrane, along with the corresponding object detection image. The scale bar represents 10 µm. The yellow insets are zoomed-in images presented as heat maps, accompanied by a calibration bar for pixel intensities. Scale bar – 2 µm (b) The morphological variants of Myomerger clusters that are detected. Scale bar – 1 µm, 100 nm (c) Objects with a having annular ring like intensity profile were also observed. Scale bar – 1 µm, 100 nm The magenta line indicated the threshold intensity that has been employed for object detection.(d) Representative level of non-specific background signal, with insets that are zoomed in. (e) Representative level of signal present in secondary negative control set of samples. Scale bar – 1 µm, 100 nm

**Figure S6**

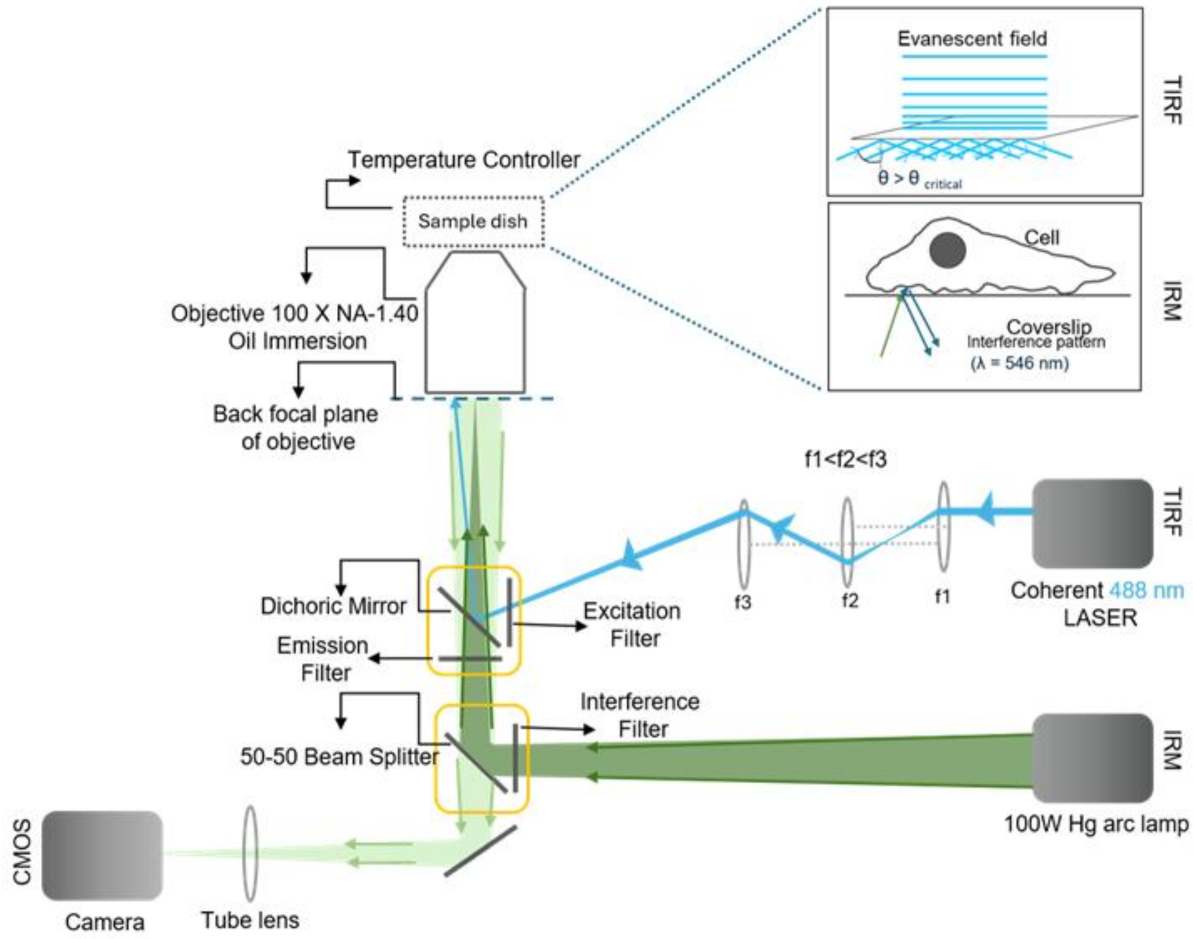


**Figure S6– Schematic optical Ray diagram of Sequential TIRF- IRM microscope system**

### Figure S7

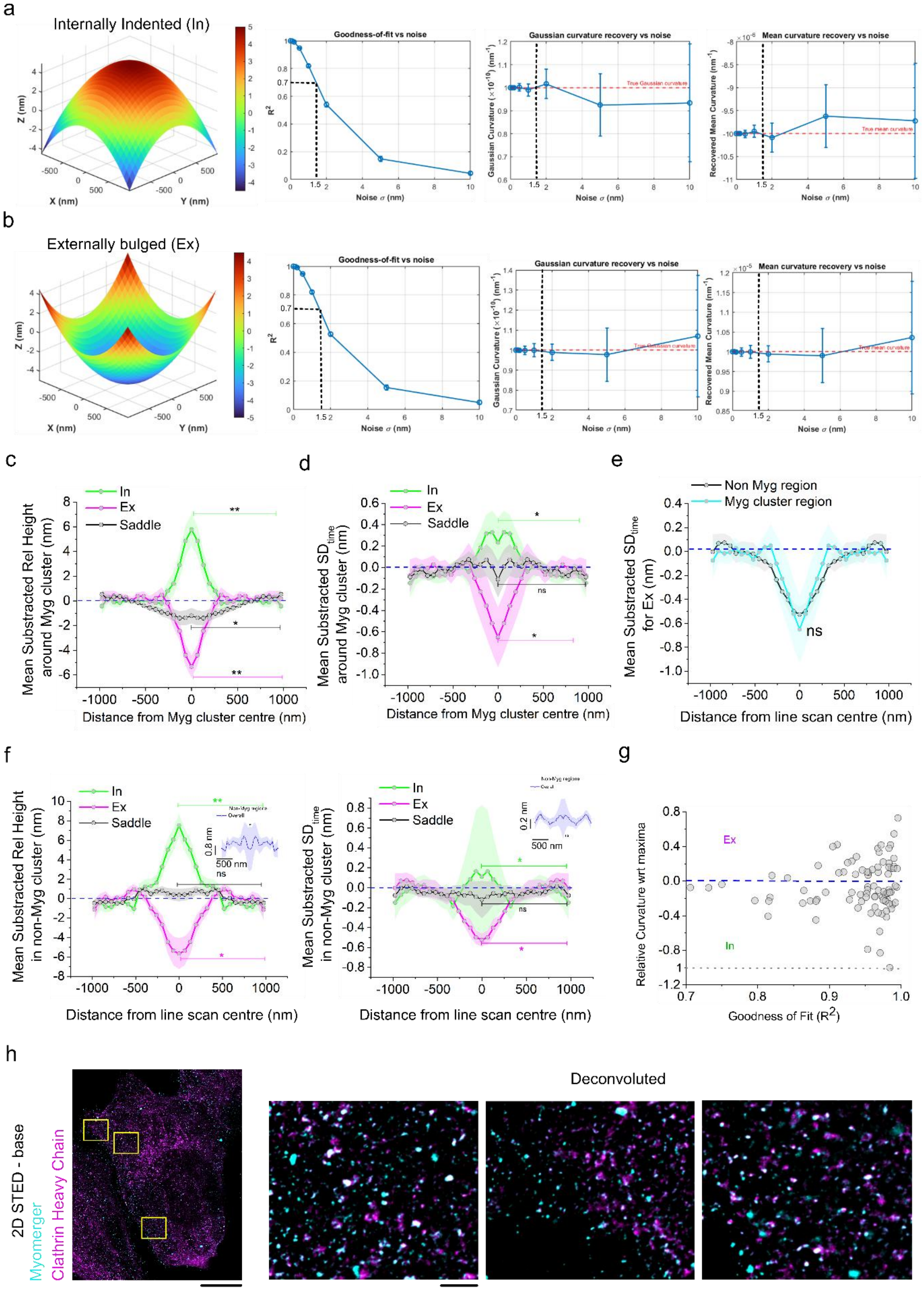

a
Internally Indented (In)
Goodness-of-fit vs noise
Gaussian curvature recovery vs noise
Mean curvature recovery vs noise
b
Externally bulged (Ex)
c
d
e
f
g
h
Deconvoluted
2D STED - base
Myomerger
Clathrin Heavy Chain
Distance from Myg cluster centre (nm)
Distance from line scan centre (nm)
Goodness of Fit ($R^2$)
Relative Curvature wrt maxima

**Figure S7– Myomerger clusters significantly alter local membrane topology**

(a,b)Simulated In (top) and Ex (bottom) topology membrane created using Quadratic equation used for fitting. The Plots mark the $R^2$ threshold of 0.7 that corresponds with a noise level of 1.5 nm. (c) Symmetric radial profiles of mean-subtracted Rel Height around cluster centroid in y axis and distance from cluster centre on x axis, for different membrane topologies. (d) Symmetric radial profiles of mean-subtracted $\Delta SD_{time}$ around cluster centroid in y axis and distance from cluster centre on x axis, for different membrane topologies (e) The comparative radial profile of mean-subtracted $\Delta SD_{time}$ of the Ex topology of Myg cluster region and non-Myg cluster region. (f) For Non Myomerger cluster non fluorescent regions from Transfected cells, (left) symmetric radial profiles of mean-subtracted Rel Height around cluster centroid in y axis and distance from cluster centre on x axis, for different membrane topologies and (right) symmetric radial profiles of mean-subtracted $\Delta SD_{time}$ around cluster centroid in y axis and distance from cluster centre on x axis, for different membrane topologies (g) Relative curvature normalized wrt maximum curvature generated at Myomerger clusters centres at In (green) and Ex (magenta) features of the membrane in y axis with corresponding goodness of fit in x-axis. The grey dotted line at 1 mark the maximum value use for normalisation. $N_{repeat}$ -3, $N_{cell}$ – 25, $N_{cluster}$ – 200, $N_{non\text{-}Myg\ cluster}$ – 985. (h) The Myomerger (cyan – IF-Abberior STAR red) and clathrin heavy chain (magenta – IF – Alexa Fluor 568) clusters co-distribution at the basal membrane of 24 hr DM treated C2C12 cells captured by 2D STED. The yellow rectangular insets are zoomed in. Scale bar 5 µm, 500 nm. * $p < 0.05$, ** $p < 0.001$, ns $p > 0.05$. The Mann-Whitney U-test was used for analysis. For all line plots, the median line profile with s.e.m as the shaded region is depicted. For all box plots, error is SD.

**Figure S8**

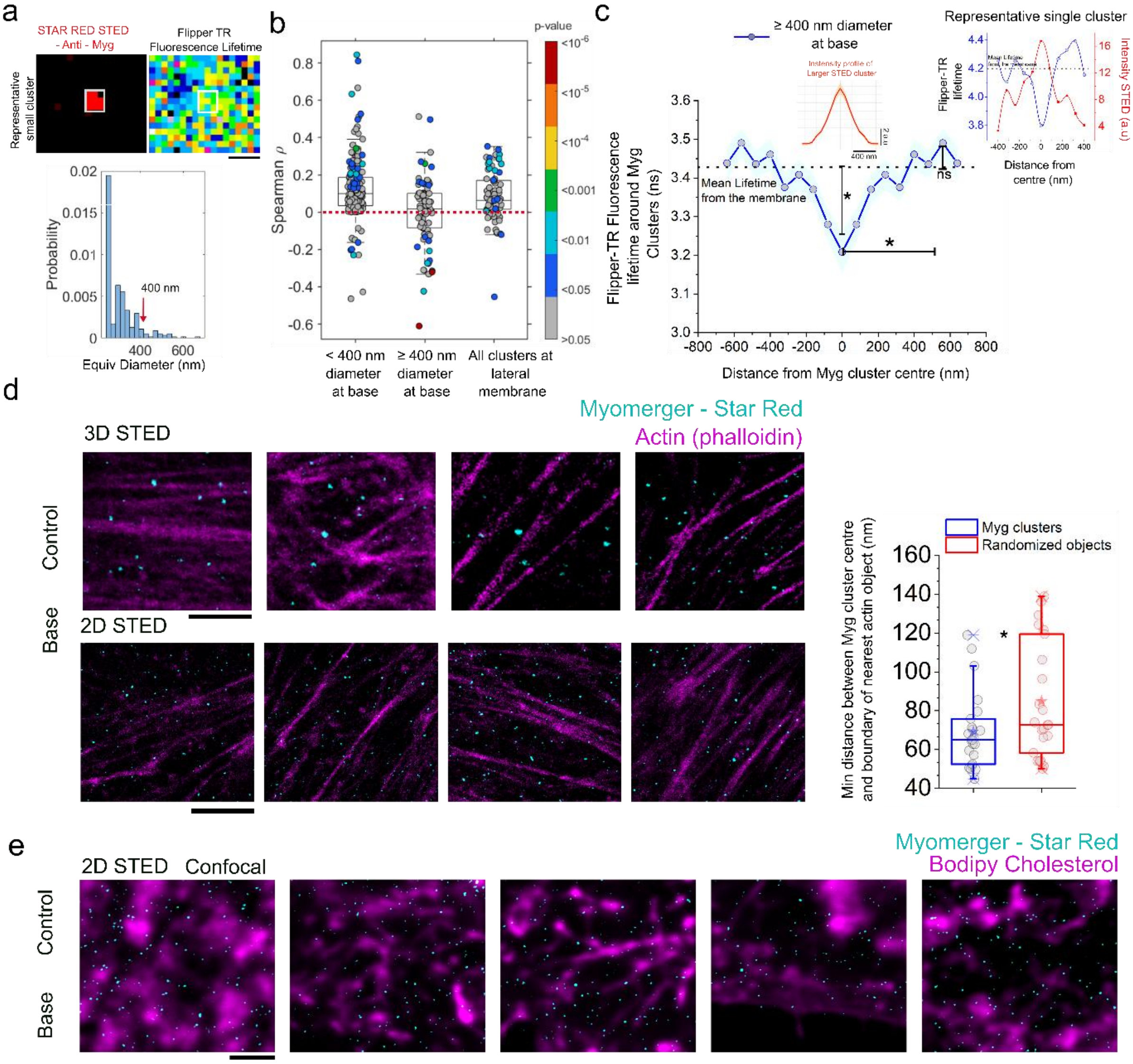

a
STAR RED STED - Anti - Myg
Flipper TR Fluorescence Lifetime
Representative small cluster
Probability
400 nm
Equiv Diameter (nm)
b
p-value
Spearman ρ
< 400 nm diameter at base
≥ 400 nm diameter at base
All clusters at lateral membrane
c
≥ 400 nm diameter at base
Representative single cluster
Mean Lifetime from the membrane
Flipper-TR Fluorescence lifetime around Myg Clusters (ns)
Distance from Myg cluster centre (nm)
Flipper-TR lifetime
Intensity STED (a.u)
Distance from centre (nm)
ns
d
3D STED
2D STED
Control
Base
Myomerger - Star Red
Actin (phalloidin)
Myg clusters
Randomized objects
Min distance between Myg cluster centre and boundary of nearest actin object (nm)
e
2D STED
Confocal
Myomerger - Star Red
Bodipy Cholesterol

**Figure S8– Smaller pool of Myomerger cluster dampens membrane fluctuation at *the centre,* and larger ones are mostly association invaginations**

(a) (top)The representative patch centred at the Myomerger cluster centroid, such that in an 8x8 region around the centroid of the Myomerger cluster, the Spearman correlation coefficient (ρ) between pixel-wise cluster intensity and FlipperTR Fluorescence Lifetime is calculated. Scale bar – 400 nm. (bottom) The normalised probability distribution (y axis) of Myomerger cluster diameters (Equivalent Diameter -x axis) (in nanometers) obtained from object detection. Red arrow marks the diameter value at which the classification of smaller vs larger clusters is done (b) The Spearman ρ value of the three classes of clusters < 400 nm, ≥ 400 nm and lateral membrane cluster, with colour code signifying the p value of correlation.(c)The symmetric profile of Flipper-TR lifetime from the basal membrane centred around the larger Myomerger cluster (≥ 400 nm in diameter). The inset shows the average radial profile of fluorescence intensity of Myomerger clusters. 0 in x-axis marks the centroid of cluster. Dotted black line marks the mean lifetime of the basal membrane. (d) Representative section from 3D and 2D STED imaging of the C2C12 cell basal membrane, taken 24 hours after DM administration. Fixed cells are immunolabeled for endogenous Myomerger (Abberior STAR Red – cyan LUT) and F-actin (Alexa Fluor 568 Phalloidin – magenta LUT). Scale bar 1 µm. The comparative study of minimum distance of real Myomerger cluster centre from the boundary of the nearest cholesterol object versus the randomised Myomerger cluster centroid positions. (e) Representative section from 2D STED - confocal imaging of the C2C12 cell basal membrane, taken 24 hours after DM administration. Fixed cells are immunolabeled for endogenous Myomerger (Abberior STAR Red - 2D STED – cyan LUT) and cholesterol (Bodipy cholesterol – confocal -magenta LUT). Scale bar 1 µm. $N_{repeat}$ -2, $N_{cell}$ – 23 (F-actin-Myg coloc); 25 (chol-Myg coloc). The Mann-Whitney U-test was used for analysis. * $p < 0.05$, ** $p < 0.001$, ns $p > 0.05$. For all line plots, the median line profile with s.e.m as the shaded region is depicted. For all box plots, error is SD.

#### Figure S9

**Figure S9 – Video of a patch of membrane that has net average IN topology for 100s.**

(a) A time-lapse series of the representative 31x31 membrane patch around Myomerger cluster centroid (marked by red solid sphere). 5x5 fitted surface in magenta that marks the local In curvature and its temporal fluctuation for 200 frames taken at interval of 10 frames of the 2048 frame IRM time lapse captured. X axis marks the distance from centre of cluster; the z-axis marks the height fluctuation. The colour bar marks the relative height.

#### Figure S10

**Figure S10 – Video of a patch of membrane that has net average EX topology for 100s.**

(a) A time-lapse series of the representative 31x31 membrane patch around Myomerger cluster centroid (marked by red solid sphere). 5x5 fitted surface in magenta that marks the local Ex curvature and its temporal fluctuation for 200 frames taken at interval of 10 frames of the 2048 frame IRM time lapse captured. X axis marks the distance from centre of cluster; the z-axis marks the height fluctuation. The colour bar marks the relative height.
